# Network Pharmacology-Guided Discovery of Fungal Autophagy Modulators for Tauopathies: Structural and Proteomic Evidence

**DOI:** 10.64898/2026.07.24.740367

**Authors:** Angel Ramon Torres Mc Cook, Camila Belen Mimura, Lautaro Damian Alvarez, Ana Clara Liberman

## Abstract

Autophagic clearance of hyperphosphorylated tau is impaired in tauopathies, leading to the progressive accumulation of toxic tau species. Fungal metabolites provide a rich yet largely untapped source of neuroactive molecules with therapeutic potential. Here, we investigated metabolites from Lion’s Mane (*Hericium erinaceus*), Magic Mushrooms (*Psilocybe* spp.), and Ergot fungi (*Claviceps* spp.) using a computational drug-discovery workflow. We characterised their structural diversity, predicted blood-brain barrier permeability and toxicological properties, and integrated network pharmacology with protein-protein interaction and functional enrichment analyses to identify autophagy-related targets. Peroxisome proliferator-activated receptor gamma (PPARG), glycogen synthase kinase-3 beta (GSK3B), and casein kinase 2 alpha 1 (CSNK2A1) emerged as the three most promising candidates, given their complementary roles linking autophagy and tau pathology and their attractiveness as targets for multi-target drug discovery. Their interactions with fungal metabolites were evaluated by molecular docking, Molecular Mechanics/Generalized Born Surface Area (MM/GBSA) rescoring, molecular dynamics simulations, complemented by machine-learning quantitative structure-activity relationship (QSAR) modelling as an additional, ligand-based line of evidence. Molecular dynamics and MM/GBSA analyses confirmed stable, target-specific binding for Corallocin A and Erinacerin M (PPARG), Chaetopyranin and Ergocryptine (GSK3B), and Hericioic Acid D and Isohericerin (CSNK2A1), alongside Emodin, a reference compound with previously reported activity against all three targets. QSAR predictions were informative primarily for the PPARG candidates, which fell within the model’s applicability domain; predictions for the remaining candidates fell outside their respective models’ applicability domains and were therefore not interpretable as evidence for or against their prioritisation. Reanalysis of an independent hippocampal proteomic dataset from Alzheimer’s disease patients showed CSNK2A1 protein levels to be significantly altered in the CA3 subfield, providing an additional, correlative line of support for this target; PPARG and GSK3B showed no significant changes in protein abundance, which does not preclude their functional involvement given their extensive post-translational regulation. Overall, these findings identify fungal metabolites as promising multi-target autophagy modulator candidates and provide a systematic computational strategy for prioritising them for experimental validation in tauopathies.

## 1. Introduction

Tauopathies constitute a group of neurodegenerative disorders that includes Alzheimer’s disease (AD) (Frost et al., 2025). They are characterized by the intracellular accumulation of hyperphosphorylated tau protein (Rawat et al., 2022), which progressively disrupts neuronal function and leads to neurodegeneration. As life expectancy increases worldwide, the clinical and socioeconomic burden associated with these disorders continues to grow, making this group of diseases one of the major challenges in modern medicine (Cummings et al., 2014). In 2019, dementia associated with AD and related tauopathies affected an estimated 57.4 million people worldwide, and this number is projected to reach 152.8 million by 2050 (Nichols et al., 2022). Despite this, currently available treatments remain largely symptomatic, and therapies capable of slowing or preventing the molecular processes that lead to neurodegeneration are still unavailable (Cummings et al., 2014; Harris et al., 2025).

A central feature of tauopathies is the loss of intracellular protein quality-control mechanisms that are primarily responsible for eliminating pathological tau. Macroautophagy (hereafter referred to as autophagy) constitutes the main pathway responsible for tau clearance (Liu et al., 2022). Indeed, it has been reported that post-mortem brain tissue from patients with AD and other tauopathies shows evidence of autophagy-lysosomal system dysfunction, including the accumulation of autophagic vesicles and impaired lysosomal maturation (Piras et al., 2016). Importantly, restoration of autophagic activity in patient-derived neurons reduces the levels of phosphorylated and insoluble tau, supporting autophagy as a promising intervention strategy for these neurodegenerative diseases (Menzies et al., 2015; Djajadikerta et al., 2020; Silva et al., 2020). In this context, natural products represent a relatively unexplored source of such alternatives.

Fungi are a rich and still underexplored source of secondary metabolites with highly diverse chemical structures and activity in the central nervous system (Wu et al., 2022a). Among fungi with neuroactive properties, three groups stand out as promising for the discovery of autophagy modulators. Lion’s Mane mushroom (*Hericium erinaceus*) produces hericenones and erinacines, metabolites that stimulate the synthesis of nerve growth factor (NGF) and promote neurotrophic and neuroprotective responses in various neuronal models (Lai et al., 2013). Hallucinogenic mushrooms (*Psilocybe* spp.) produce psilocybin and other related indole alkaloids that, after conversion into their active metabolites, activate serotonergic signaling pathways involved in neuroplasticity, with sustained effects over time (Calder and Hasler, 2023). Ergot fungi (*Claviceps* spp.) synthesize ergot or ergoline alkaloids that interact with dopaminergic, serotonergic, and adrenergic receptors and have served as structural templates for the development of drugs targeting the central nervous system (Mantegani et al., 1999). Taken together, the chemical diversity and neuroactive properties of the metabolites produced by these three fungal groups make them a promising and largely unexplored source of compounds for regulating autophagy in tauopathies.

The multifactorial nature of tauopathies challenges the traditional one drug-one target paradigm, highlighting the need for therapeutic strategies capable of modulating multiple disease mechanisms simultaneously. Network pharmacology has emerged as an effective approach by integrating systems biology with computational methods to identify compounds that act on several disease-relevant targets (Li and Kar, 2025). However, most studies are still restricted to target identification followed by molecular docking analysis. Although docking is useful for predicting ligand–protein interactions, it does not provide sufficient information to estimate binding affinity. Molecular dynamics simulations, blood-brain barrier permeability prediction and toxicological assessment are frequently not included (Pereira et al., 2021). Similarly, ligand-based approaches such as QSAR modelling are incorporated only in a limited number of studies. Computational predictions are also rarely evaluated using patient-derived datasets. Integrating human omics data allows determining whether the identified targets are also dysregulated in affected tissues, thus providing independent biological support for the computational predictions.

Previous computational studies have explored the potential of fungal metabolites in the context of AD. These studies analyzed their interaction with molecular targets associated with the disease, such as GSK-3β, the NMDA receptor, and BACE-1, using in silico tools such as pharmacophore modelling, molecular docking, and molecular dynamics simulations (Iqbal et al., 2023). Similarly, network pharmacology studies performed with psilocybin-producing fungi identified monoaminergic and G protein-coupled receptors, including HTR2A, MAOA, and DRD2, as possible targets related to their neuropsychiatric effects (Murray et al., 2026). However, these studies were not designed to investigate autophagy regulation in tauopathies. In addition, most of these approaches do not incorporate other relevant parameters to evaluate the therapeutic potential of the compounds, such as blood-brain barrier permeability, toxicological profile, complementary affinity analyses through ligand-based approaches, or validation with independent patient-derived data. The integration of these tools with human biological information is necessary to improve the selection of candidates with greater therapeutic relevance.

In this study, we investigated fungal metabolites from Lion’s Mane, *Psilocybe* spp., and *Claviceps* spp. as potential multi-target modulators of autophagy in tauopathies, using an integrated computational strategy. We combined network pharmacology, protein–protein interaction analysis, and functional enrichment analysis to identify molecular targets shared between fungal metabolites, tauopathy-associated pathways, and the autophagy network. PPARG, GSK3B, and CSNK2A1 emerged from this analysis as central components of an autophagy-related subnetwork, which we then examined in greater detail.

These three proteins act on autophagy and tau pathology through different mechanisms. GSK3B phosphorylates tau protein at multiple sites associated with the aggregation process and at the same time inhibits autophagy through mTOR-dependent signalling (Lior et al., 2025).CSNK2A1 promotes tau hyperphosphorylation through the regulation of the SET/PP2A axis and is increased in the brain with AD, suggesting that it could contribute to the accumulation of pathological tau (Zhang et al., 2018). CSNK2 has also been characterised as a negative regulator of autophagy initiation in other pathological contexts, acting through phosphorylation of FLN-NHL–containing TRIM proteins to inhibit the ULK1-BECN1 complex (Hoenigsperger et al., 2023), although its specific role in autophagy regulation within tauopathies remains largely unexplored. PPARG regulates the expression of genes related to autophagy and participates in the maintenance of mitochondrial function, positioning it as a relevant regulator of cellular processes altered in tauopathies(Ahmed et al., 2019; Praharaj et al., 2024). Pharmacological activation of PPARG has also been shown to reduce tau accumulation and hyperphosphorylation in neuronal models, indicating a direct link between PPARG activity and tau pathology (Moosecker et al., 2019).

The involvement of these proteins in different cellular processes led us to evaluate a multi-target approach, rather than focusing on a single protein. For this purpose, we combined structural analyses, ligand-based predictions, and patient-derived data. We integrated network pharmacology, molecular modelling, ligand-based predictions, and human proteomic data to identify fungal metabolites with potential modulatory effects on autophagy. This approach allowed the prioritization of multi-target candidates for future experimental validation in tauopathies.

## 2. MATERIALS AND METHODS

Figure 1 shows the workflow used in this study, from the creation of the library of fungus-derived compounds to the prediction of binding affinities using QSAR models.

**Figure 1.**
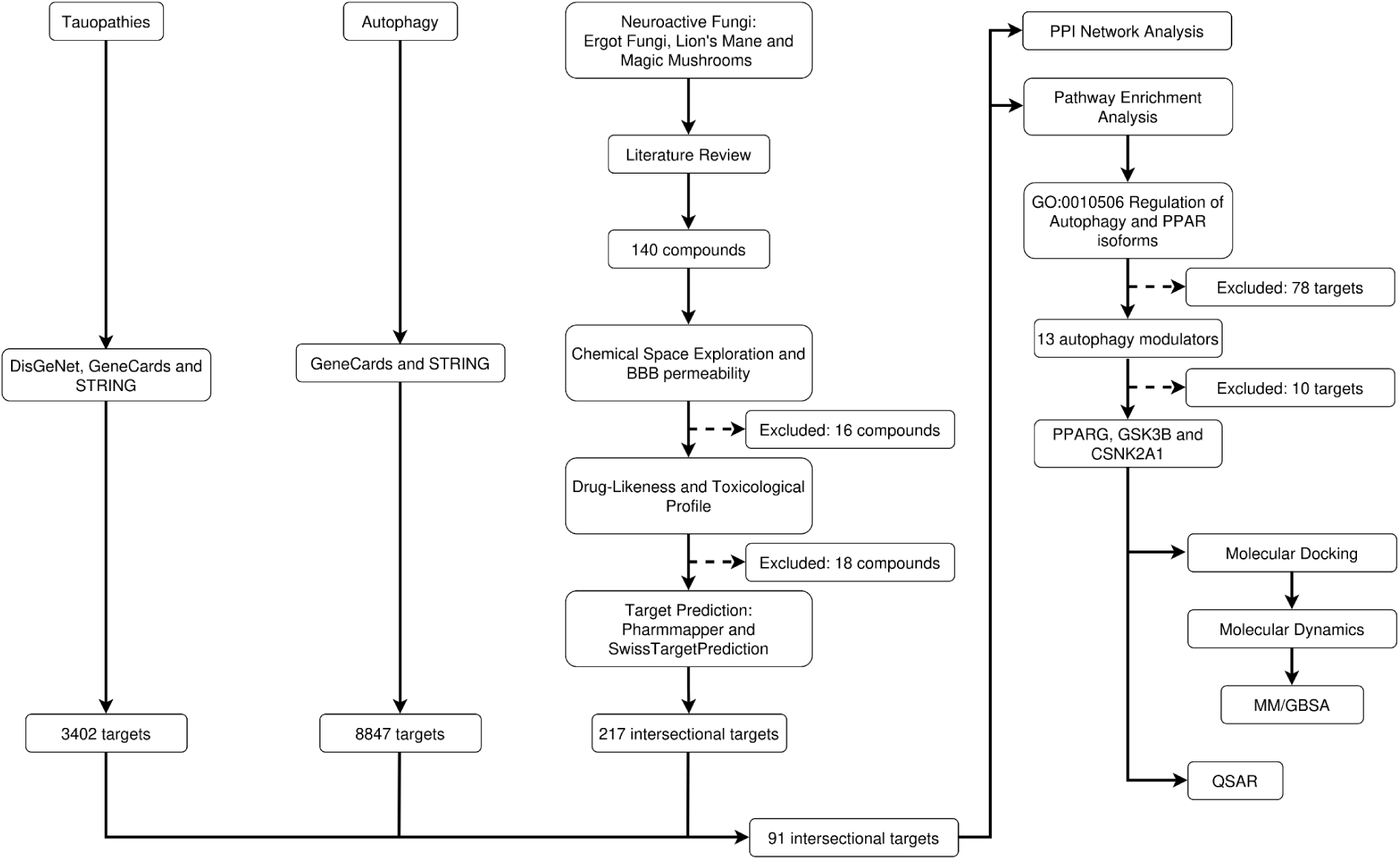
Workflow of the network pharmacology-based analysis of the mechanisms through which neuroactive fungi regulate autophagy in the context of tauopathies.

### 2.1. Construction of the fungal-derived compound library

We compiled a library of neurobioactive compounds derived from fungi, through a review of the scientific literature using PubMed and Google Scholar. For this, we focused on three taxonomically distinct fungal sources, with previously described neuroactive or neuromodulatory properties: Ergot Fungi (*Claviceps* spp.), Lion’s Mane (*Hericium erinaceus*), and Magic Mushrooms (*Psilocybe* spp.). The search included combinations of the terms “ergot alkaloids” and “neuroactive,” “*Hericium erinaceus* compounds,” and “*Psilocybe* metabolites,” without restrictions on publication date.

The chemical structure of each identified compound was obtained from the PubChem database. In the case of compounds not listed in PubChem, structures were manually constructed from the chemical information gathered from the source literature. All structures were processed with Open Babel v3.1.1(O’Boyle et al., 2011) to generate three-dimensional conformations. This ensured that 3D molecular descriptors could be calculated consistently. No duplicate compounds were found among the three fungal sources, resulting in a library of 140 compounds (Supplementary Figures 1-3).

### 2.2. Molecular descriptor calculation and chemical space analysis

The molecular descriptors of each compound were calculated using the rcdk v3.8.2 package (Guha, 2007; Voicu et al., 2020) in R v4.6.1. Since the library was composed exclusively of small non-peptide and non-protein molecules, protein-specific descriptor categories were excluded from the calculation. Next, hybrid, constitutional, topological, electronic, and geometric descriptors were calculated for each compound.

Before dimensionality reduction, we removed descriptors that contained missing values or a constant value of zero across all compounds, since such variables cannot be meaningfully standardized. Next, the remaining descriptors were centered and scaled using z-score normalization. We performed a principal component analysis (PCA) on the scaled descriptor matrix and retained the first ten principal components. Subsequently, the first two principal components were used for visualization and subsequent clustering.

We determined the optimal number of clusters by applying the elbow method to the scores of the first two principal components. This method uses the within-cluster sum of squares, implemented in the factoextra package (Kassambara and Mundt, 2026).

### 2.3. ADMET and drug-likeness prediction

#### 2.3.1. Blood-brain barrier permeability prediction

We evaluated the ability of the library of fungal-derived compounds to cross the blood-brain barrier (BBB) as a first selection criterion. We calculated the predicted logBB (pLogBB) values using LogBB_Pred, a quantitative machine learning-based model for predicting BBB permeability (Shaker et al., 2023). Compounds with a pLogBB > −1 were classified as BBB-permeable and retained for further analysis.

We evaluated differences in pLogBB among the structural clusters previously identified through a clustering analysis based on principal component analysis (PCA), using a PERMANOVA, implemented in the vegan v2.7-5 package (Oksanen et al., 2026). Subsequently, pairwise post hoc comparisons were performed among clusters using the R package RVAideMemoire v0.9-83-12 (Herve, 2025). Differences were considered statistically significant with a p-value < 0.05.

#### 2.3.2. Drug-likeness assessment: Lipinski’s Rule of Five

As a second selection criterion, compliance with Lipinski’s Rule of Five (Lipinski et al., 2001) was evaluated for the subset of compounds that passed the BBB permeability filter. Compounds that did not comply with at least two of the four rules were considered physicochemically suboptimal for absorption and oral bioavailability, and were therefore excluded from subsequent analyses.

#### 2.3.3. Toxicity prediction

The toxicity profiles of the subset of BBB-permeable compounds were obtained from ProTox 3.0 (Banerjee et al., 2024). These profiles included six categories: organ toxicity, toxicological endpoints, nuclear receptor signaling pathways, stress response pathways, molecular initiating events, and metabolism-related endpoints. The toxicity results were used solely for descriptive and comparative purposes among the different groups of chemical compounds and fungal sources.

### 2.4. Prediction of fungal-derived compound targets

The possible protein targets of each fungal-derived compound in the library were predicted using two complementary platforms: SwissTargetPrediction (Daina et al., 2019) and PharmMapper (Liu et al., 2010). In SwissTargetPrediction, the canonical SMILES notation of each compound was entered as input, the organism was set as Homo sapiens, and the standard target filtering threshold was set at a probability > 0. In PharmMapper, the three-dimensional structure of each compound was entered as input, in SDF format, the species was set as Homo sapiens, and a standard Z-score threshold ≥ 2 was applied to filter the predicted targets.

For each compound, the lists of predicted targets were standardized to official gene symbols using UniProt (The UniProt Consortium, 2025). Only the targets predicted by both SwissTargetPrediction and PharmMapper were retained for each compound. Next, duplicates derived from targets shared between different compounds were removed. This yielded a final, consolidated, and non-redundant set of possible protein targets associated with the fungal-derived compounds. This set was used for subsequent network pharmacology analyses.

### 2.5. Collection of tauopathy-and autophagy-associated targets

Targets associated with tauopathy were obtained from three databases: DisGeNet (Hu et al., 2025), GeneCards, and STRING (Szklarczyk et al., 2025), using the search term “tauopathy”. Targets related to autophagy were obtained from GeneCards and STRING using the search term “autophagy”. In both cases, genes that do not encode proteins were excluded from the GeneCards results. All searches were performed in October 2025.

The obtained target lists were standardized using the official UniProt gene symbols for each category. Then, they were merged and duplicates were removed. This yielded two final, consolidated, and non-redundant sets of targets associated with tauopathy and autophagy, respectively.

### 2.6. Target overlap and intersection analysis

To identify targets shared among fungal-derived compounds, tauopathies, and autophagy, we determined the intersection of the targets associated with each of these three categories. The targets that overlapped among the three sets were retained for downstream network pharmacology analyses.

To further characterize this overlapping target set, its distribution among the three fungal sources was examined. First, the predicted target list of each source was subset to include only the targets present in the above-identified overlapping set. Then, the intersection of these three subsets was determined to identify the targets shared across the three sources.

### 2.7. Gene Ontology and KEGG pathway enrichment analysis

The targets that overlapped between the fungal-derived, tauopathy-associated, and autophagy-related targets were annotated with their corresponding Entrez gene identifiers using the genekitr package v1.2.8 (Liu and Li, 2023) with the organism set to human.

We performed Gene Ontology (GO) enrichment analyses separately for the biological process (BP), cellular component (CC), and molecular function (MF) categories, with the clusterProfiler package v4.20.0 (Yu et al., 2012). For each category, we applied a p-value cutoff of 0.05, a q-value cutoff of 0.05, a minimum gene set size of 10, and a maximum gene set size of 500.

A KEGG pathway enrichment analysis was performed using clusterProfiler. The organism was set to Homo sapiens, and the same significance thresholds and gene set size limits were applied as in the GO enrichment analyses. For all four enrichment analyses, only terms and pathways with an p_adj_ < 0.05 were considered statistically significant.

### 2.8. Protein-protein interaction network construction

The overlapping targets of the fungal-derived, tauopathy-associated, and autophagy-related compounds were imported into the STRING database for constructing a protein-protein interaction (PPI) network. The species was set to Homo sapiens and the minimum interaction score was set to a high-confidence threshold of 0.7. The resulting PPI data were exported from STRING as a TSV file and imported into Cytoscape v3.10.4(Shannon et al., 2003) for network visualization and topological analysis.

We used the CytoNCA plug-in (Tang et al., 2015) in Cytoscape to calculate three topological parameters for each node in the network: degree centrality (DC), betweenness centrality (BC), and closeness centrality (CC).

### 2.9. Molecular docking

We performed molecular docking assays using AutoDock Vina v1.2.0 (Trott and Olson, 2010) to evaluate the binding interactions between 40 prioritized, fungal-derived compounds and their corresponding predicted targets among PPARG, GSK3B, and CSNK2A1. Emodin was included among these 40 compounds as an internal reference, given its previously reported activity against the three targets (Yamada et al., 2005; Gebhardt et al., 2010a; Li et al., 2025a). This allowed us to use its docking results as a benchmark for the remaining candidate compounds.

The crystallographic structures of the three target proteins were retrieved from the Protein Data Bank (PDB): the ligand-binding domain of PPARG (PDB ID: 1PRG), GSK3B (PDB ID: 6Y9S), and CSNK2A1 (PDB ID: 3Q9X). Each structure was prepared using VMD v1.9.4a57 (Humphrey et al., 1996), and then converted to PDBQT format using AutoDockTools (El-Hachem et al., 2017).

The 40 candidate compound three-dimensional structures were first converted from SDF to PDB format using OpenBabel, and subsequently converted to PDBQT format using AutoDockTools.

For each target, a grid box with 0.375 Å spacing was centered on the ligand-binding pocket of PPARG and the ATP-binding sites of GSK3B and CSNK2A1. Grid box dimensions were set for each target according to its binding site: 17 × 20 × 20 points for PPARG, 16 × 20 × 16 points for GSK3B, and 16 × 16 × 20 points for CSNK2A1. During docking, ligands were treated as fully flexible with respect to their rotatable bonds, while the receptor was kept rigid.

Docking calculations were performed with an exhaustiveness value of 8, generating nine binding modes per ligand within an energy range of 3 kcal/mol. Each compound was docked only against targets with which it had previously shown a predicted interaction. The pose with the lowest predicted binding affinity, as reported by AutoDock Vina, was selected for each compound-target complex and used as the starting conformation for molecular dynamics simulations.

### 2.10. Molecular dynamics simulations

The initial protein-ligand complexes for PPARG, GSK3B, and CSNK2A1 were constructed using the corresponding PDB coordinates and docking solutions obtained for the prioritized fungal-derived compounds. For GSK3B, we modeled an unresolved short loop in the crystallographic structure using Modeller v10.5 (Webb and Sali, 2016).

All molecular dynamics (MD) simulations were performed with AMBER22 (Case et al., 2023). Protein parameters were taken from the ff14SB force field, and ligand parameters were assigned using the General AMBER Force Field (GAFF). Atomic charges were derived as RESP charges at the HF/6-31G** level. Each system was solvated in an octahedral TIP3P water box. Then, each system was subjected to a relaxation and equilibration protocol consisting of solvent minimization, staged heating, and a multi-step NVT-NPT equilibration scheme prior to production.

Solvent minimization was performed using conjugate gradient minimization, with harmonic positional restraints of 100 kcal·mol⁻¹·Å⁻² applied to the solute’s heavy atoms. The systems were then heated from 100 K to 298 K over a 1.0 ns NVT segment (1 fs timestep), while maintaining strong positional restraints (100 kcal·mol⁻¹·Å⁻²) on the protein and ligand atoms. Subsequent equilibration consisted of a series of 1-ns NPT segments (1-fs timestep) with progressively relaxed restraints on the solute, followed by a final unrestrained NPT segment. The SHAKE algorithm constrained bonds involving hydrogen atoms and allowed the above-described timesteps. A nonbonded interaction cutoff of 8.0 Å was used throughout. Pressure (1 atm) was maintained using AMBER’s Monte Carlo barostat under isotropic periodic boundary conditions. Temperature was regulated at 298 K using the Langevin thermostat.

Production simulations were performed in the NPT ensemble with a 2 fs time step. These simulations continued directly from the final equilibrated restart file, using the same thermostat and barostat settings described above. For each prioritized protein-ligand complex, production was carried out in triplicate, with three independent 300 ns replicates simulated per complex.

### 2.11. Molecular dynamics trajectory analysis

#### 2.11.1. Trajectory clustering

For each protein-ligand complex, the three 300 ns production replicates were concatenated and subjected to conformational clustering using the density-based clustering algorithm DBSCAN, as implemented in cpptraj (Roe and Cheatham, 2013). Clustering was performed on the RMSD calculated over the ligand atoms, with a minimum cluster size of 25 points and a distance threshold of 0.9 Å. To reduce computational cost, a sieving factor of 5 was applied during the clustering step. All original frames were subsequently reassigned to their corresponding cluster using the sieveToFrame command. For each complex, the representative conformation of the most populated cluster was extracted directly from the trajectory as the frame closest to the cluster centroid.

#### 2.11.2. Pairwise RMSD analysis

Pairwise RMSD analysis was performed independently for each protein-ligand complex and each of the three production replicas using the MDTraj Python library (McGibbon et al., 2015). Each trajectory was superimposed onto the protein backbone atoms using the first frame of the same trajectory as the reference structure. The Cartesian coordinates of the ligand’s heavy (non-hydrogen) atoms were extracted from the superposed trajectory. A pairwise RMSD matrix was constructed by comparing these coordinates between all pairs of frames within the same protein-anchored reference frame. To reduce computational cost, trajectories were subsampled to a maximum of 1,500 frames, using a stride calculated automatically based on the total number of frames in each trajectory.

#### 2.11.3. Principal component analysis of ligand motion

For each protein-ligand complex, we performed PCA independently for each of the three production replicas. Each trajectory was superimposed onto the protein backbone atoms using the first frame of the same trajectory as the reference structure. The Cartesian coordinates of the ligand’s heavy (non-hydrogen) atoms were then extracted from the superimposed trajectory and used as input variables for PCA. We performed PCA using the implementation from the scikit-learn library (Pedregosa et al., 2011) and retained the first three principal components for each replica, along with their corresponding explained variance ratios.

For each complex, the trajectory was projected onto the first two principal components to visualize the conformational space sampled by the ligand throughout the simulation.

### 2.12. MM/GBSA binding free energy calculations

Binding free energies (ΔG_bind_) were estimated using the Molecular Mechanics/Generalized Born Surface Area (MM/GBSA) approach implemented in the MMPBSA.py module of AmberTools22 (Case et al., 2023). The analysis was performed independently on each of the three production replicas for each protein-ligand complex. From each 300 ns trajectory, cpptraj extracted frames spanning the entire simulation with a stride of 3. This yielded a subset of 1,000 frames per replica for the MM/GBSA calculation. The receptor, ligand, and complex topology files were generated from the corresponding complex topology using the ante-MMPBSA.py utility, and the system was stripped of water molecules and counterions.

MM/GBSA calculations were performed using the generalized Born model with the OBC2 parameterization and an implicit salt concentration of 0.150 M. The energetic contributions were decomposed into electrostatic and van der Waals terms. The overall ΔG_bind_ was calculated as the sum of these terms plus the polar and nonpolar solvation contributions.

### 2.13. Complementary ligand-based QSAR modelling

As a complementary, ligand-based line of evidence to the structure-based docking and MD/MM-GBSA analyses, QSAR models were developed for each target using training data obtained from BindingDB (Liu et al., 2007), which included Kd values for PPARG, as well as Ki values for GSK3B and CSNK2A1. For a subset of training compounds lacking defined three-dimensional structures, structures were generated using Open Babel v.3.1.1. Then, molecular descriptors (hybrid, constitutional, topological, electronic, and geometrical) were calculated for each set of training compounds using the rcdk package v3.8.2 in R.

Binding affinity values were converted from nanomolar (nM) to molar (M) and then transformed into their negative logarithms (pKd or pKi) to create a more suitable scale for model training. For each target, descriptors consisting entirely of missing values, those with a constant value of zero across all compounds, and those with only a single unique value across the entire training set were removed. Duplicate compounds within each training set were identified, and only those with an experimental discrepancy in pKd or pKi below a target-specific threshold were retained and had their affinity values averaged. A discrepancy threshold of ≤ 0.5 log units was applied to PPARG, and ≤ 0.1 log units to GSK3B and CSNK2A1.

We removed descriptors with near-zero variance using the caret package v7.0-1(Kuhn, 2008) with the default frequency-ratio and unique-value cutoffs. Missing values in the remaining descriptors were imputed using median imputation. Then, highly correlated descriptors were removed with a correlation cutoff of 0.90. Only one descriptor was retained from each group of correlated variables. Any remaining linearly dependent combinations of descriptors were subsequently removed.

Feature selection was performed using the Least Absolute Shrinkage and Selection Operator (LASSO) regression method. This method was implemented via the glmnet package v5.0 (Friedman et al., 2010). The method used a Gaussian family and 10-fold cross-validation. Only descriptors with non-zero coefficients were retained for model training for each target. Descriptors selected at lambda.1se were used for PPARG, and descriptors selected at lambda.min were used for GSK3B and CSNK2A1.

For each target, we split the resulting dataset into a training (80%) and testing (20%) subset. We simultaneously trained six regression algorithms, Elastic Net, Random Forest, Support Vector Regression with a radial kernel, Cubist, Gradient Boosting Machine, and Partial Least Squares, on the training subset using the caretEnsemble package v4.0.2(Deane-Mayer et al., 2026). Model training was performed using a repeated 10-fold cross-validation scheme with automatic hyperparameter tuning over 10 combinations per algorithm and root mean squared error (RMSE) as the optimization metric.

### 2.14. Model validation

#### 2.14.1. Y-randomization

A Y-randomization test was performed on the final QSAR model for each target. Using the optimal hyperparameters previously identified, the response variable in the training set was randomly permuted 1,000 times while keeping the descriptor matrix unchanged. For each permutation, a model was retrained with the same hyperparameters and cross-validation scheme. Its cross-validated Q² was then calculated to generate a null distribution. Then, the cross-validated Q² of the original, non-permuted model was compared against this null distribution.

#### 2.14.2. Applicability domain analysis

We evaluated the applicability domain of each final QSAR model using the Williams plot approach. For each target, the leverage (h) of each training and testing compound was calculated from the descriptor matrix (X) of the training set as follows:

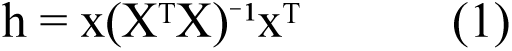

where x corresponds to the descriptor vector of a given compound.

The critical leverage threshold (h*) was defined as follows:

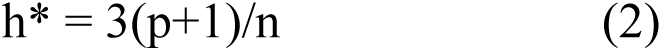

where p is the number of descriptors retained in the final model, and n is the number of compounds in the training set. Compounds with a leverage value exceeding h* were considered structurally influential outliers relative to the training chemical space.

The residuals between the experimental and predicted pKd or pKi values were calculated for both the training and testing sets, then standardized (z-scored). Compounds with an absolute standardized residual greater than 3 were considered outliers in the response.

### 2.15. Application of QSAR models to fungal-derived compounds

The three final QSAR models, developed and internally cross-validated as described above, were then applied to estimate the binding affinity of a subset of previously prioritized fungal-derived compounds, with predictions interpreted according to each model’s applicability domain.

The same molecular descriptors that were previously calculated using the rcdk package were used for each compound. Only the subset of descriptors that was retained in the final model for each target following LASSO-based feature selection was selected. Binding affinities were predicted using the algorithm selected as the final model for each target.

To determine if the evaluated fungal compounds fell within the applicability domain of each model, we calculated their leverage values using the corresponding training set’s descriptor matrix. We used the same equation to define the applicability domain of the training and testing compounds. The obtained leverage values for each compound were then compared against the previously calculated h* for each model. Since these compounds lacked experimental pKd or pKi values, applicability domain classification was based exclusively on the leverage criterion without assessing standardized residuals.

### 2.16. Reanalysis of human hippocampal proteomic data

As an independent, complementary analysis not depicted in Figure 1, quantitative proteomic data from human hippocampal subfields were obtained from (Gao et al., 2022). They characterized proteomic changes in patients with AD across hippocampal subfields using two independent TMT-based experiments (10-and 6-plex). The corresponding mass spectrometry proteomics data were deposited in the ProteomeXchange Consortium (ProteomeCentral Data and Tools, n.d.) under dataset identifier PXD027380, and were retrieved for reanalysis in the present study.

Briefly, the mass spectrometry data from the two TMT-labeled experiments were reanalyzed using the same search and quantification criteria described in the original study. MaxQuant version 2.8.0.0 (Cox and Mann, 2008) was used instead of Proteome Discoverer 2.2. The raw spectra were searched using the built-in Andromeda search engine against the reviewed human protein FASTA database, which was downloaded from UniProt in June 2026. The search allowed for a maximum of two missed cleavages, a precursor mass tolerance of 20 ppm, and a fragment mass tolerance of 0.02 Da. Carbamidomethylation of cysteine and TMT 6-plex or 10-plex labeling of peptide N-termini were set as fixed modifications. TMT 6-plex or 10-plex labeling of lysine, oxidation of methionine, phosphorylation of serine, threonine, and tyrosine, as well as deamidation of asparagine, were set as variable modifications. Protein identification was considered valid when at least one peptide was detected with an FDR below 1%. All other parameters were kept at their default values.

Peptides were quantified based on unique peptide ratios derived from TMT reporter ion signals detected by MS/MS. Quantification was considered reliable for proteins with a score greater than 10 and at least one unique peptide. The differential expression threshold was defined using the 95% prediction interval. The threshold was set to log2(ratio) ≥0.42 or ≤-0.58 for 6-plex-labeled samples and log2(ratio) ≥0.32 or ≤-0.32 for 10-plex-labeled samples. Differential expression and downstream functional enrichment analyses were performed independently for each hippocampal subfield available in the dataset. Given the exploratory nature of this region-by-region comparison, no additional correction for the number of subfields tested was applied; consequently, region-specific findings, including any subfield-level enrichment result, should be interpreted as hypothesis-generating rather than confirmatory.

## 3. RESULTS

### 3.1. Chemical space exploration, drug-likeness and ADMET prediction

To investigate the neuroprotective potential of fungal-derived compounds against tauopathy-and autophagy-related targets, we implemented an integrative computational strategy combining chemoinformatics, network pharmacology, molecular docking and dynamics simulations, binding free energy calculations, and QSAR modeling, as summarized in Figure 1. As a first step, we assembled a library of 140 neurobioactive compounds from three taxonomically distinct fungal sources with previously reported neuroactive or neuromodulatory properties: Ergot fungi, Lion’s Mane, and Magic Mushrooms. A principal component analysis (PCA) based on molecular descriptors was performed to characterize the chemical space of the library of neurobioactive fungal compounds (Figure 2A). The first two principal components explained 55.12% of the total variance, and a cluster analysis of these components revealed four distinct chemical clusters. Compounds from the three fungal sources were grouped by clusters 1, 2, and 3. This indicates that metabolites from taxonomically distinct sources may share physicochemical properties and occupy overlapping regions of chemical space. In contrast, cluster 4 consisted exclusively of Lion’s Mane compounds, which were distributed in a clearly separated region, suggesting that this subset possesses a distinctive physicochemical identity within the library.

**Figure 2.**
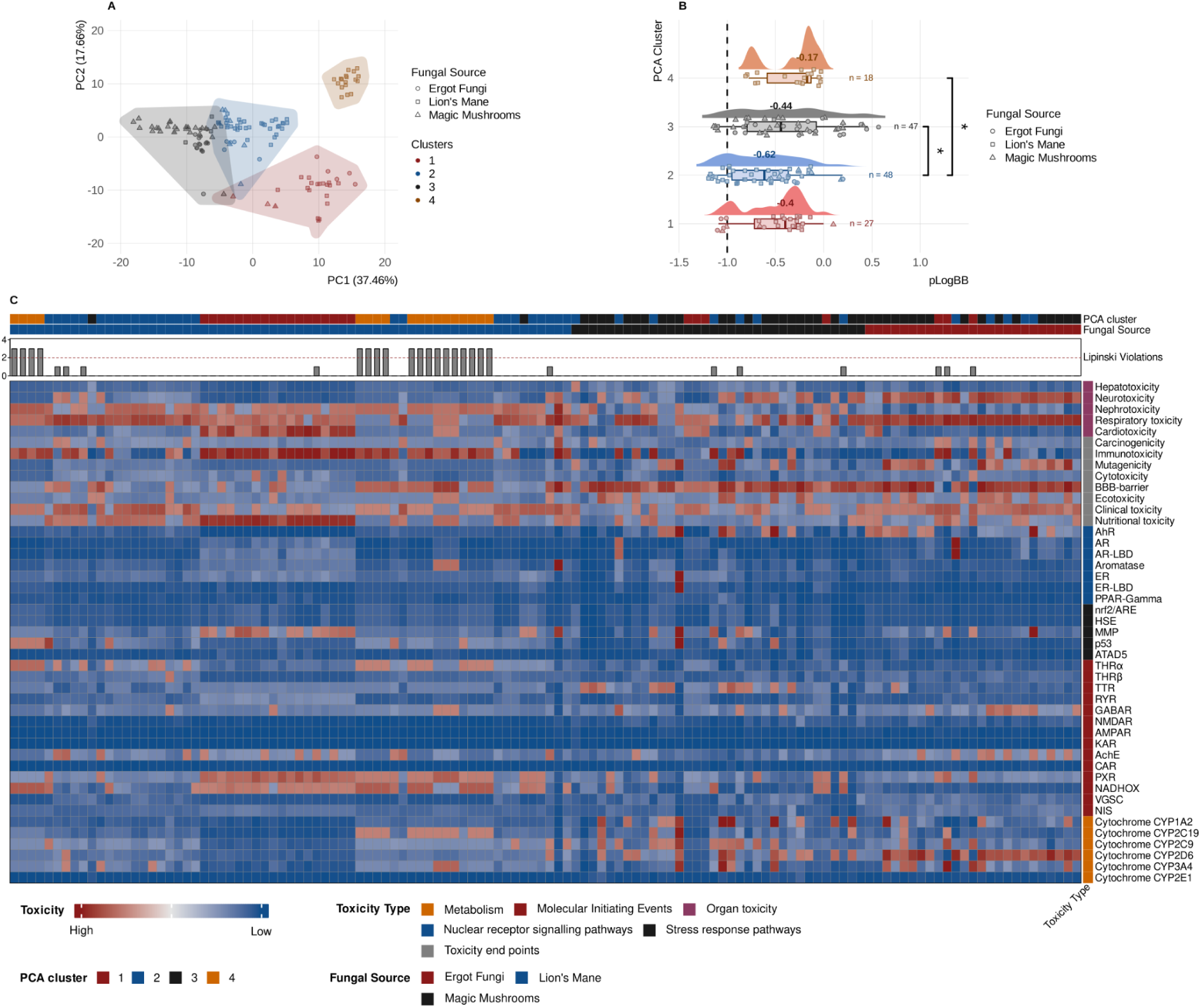
Exploration of the chemical space, predicted blood-brain barrier permeability and toxicological profile of fungal-derived compounds. (A) Principal component analysis (PCA) based on the molecular descriptors of the compounds. Individual compounds are represented by points; the shape of the symbols indicates the fungal group of origin and the colours correspond to the clusters identified by cluster analysis. The polygons delineate the distribution of each cluster within the chemical space. (B) Distribution of the predicted blood-brain barrier permeability (pLogBB) values for each chemical cluster. Each point represents an individual compound; the boxplots indicate the median (central line) and interquartile range, and the number above each distribution corresponds to the median pLogBB value of that cluster. The dashed vertical line marks the pLogBB =-1 threshold used to classify compounds as BBB-permeant. Asterisks indicate statistically significant pairwise differences in pLogBB between clusters (p < 0.05). (C) Heatmap of the predicted toxicological profile for each compound. The columns represent individual compounds, while the rows correspond to different toxicological endpoints, nuclear receptors, stress response pathways, and metabolic enzymes. Colour intensity indicates the predicted level of toxicity (red: higher toxicity; blue: lower toxicity). The top annotations show the cluster assigned by PCA, the fungal origin of each compound, and the number of violations of Lipinski’s rule.

Since tauopathies are neurodegenerative diseases characterized by the pathological accumulation of tau in neurons of the central nervous system, any candidate compound intended to modulate autophagy for therapeutic purposes must be able to cross the blood-brain barrier (BBB) to reach its site of action. Therefore, the predicted BBB permeability (pLogBB) of the 140 compounds in the library was used as the first selection criterion, comparing their distribution across the four chemical clusters identified by PCA (Figure 2B). PERMANOVA analysis revealed significant differences in pLogBB values across clusters (F = 4.62, p < 0.01). Pairwise post hoc comparisons confirmed significant differences between clusters 2 and 3 and between clusters 2 and 4 (p < 0.05), while the remaining comparisons did not reach statistical significance. The median pLogBB values were-0.40,-0.62,-0.44, and-0.17 for clusters 1, 2, 3, and 4, respectively. Cluster 4 exhibited the highest predicted BBB permeability, and cluster 2 exhibited the lowest. Using the conventional threshold of pLogBB >-1 to classify a compound as BBB-permeable, 124 compounds met this criterion. The remaining 16 compounds were classified as non-permeable and excluded from subsequent analyses, reducing the library to 124 compounds.

Of the 124 compounds that passed the BBB permeability filter, compliance with Lipinski’s rules was evaluated as a criterion for “drug-likeness” (Figure 2C). Compounds that failed to meet at least two of the four rules (n=18) were excluded from subsequent analyses since a greater number of violations is associated with suboptimal physicochemical properties for absorption and oral bioavailability. Following this screening, the library was reduced to 106 compounds for subsequent analyses.

Unlike BBB permeability and compliance with Lipinski’s rules, predicted toxicity (Figure 2C) was not an exclusion criterion. The reliability of QSTR models for toxicity prediction depends heavily on the specific chemical space covered by their training data. Although platforms like ProTox 3.0 incorporate diverse databases, fungal metabolites have highly complex and unique structural frameworks that often fall outside standard training sets. This structural divergence can decrease the precision and confidence scores of predictions for these natural products. For this reason, the toxicity profile was used for descriptive and comparative purposes across chemical clusters rather than as a binary inclusion/exclusion filter.

The predicted toxicological profile revealed heterogeneous patterns depending on the evaluated category (Figure 2C). Organ toxicity endpoints showed high toxicity predictions across the entire library, regardless of the chemical cluster or fungal origin of the compounds. This widespread pattern suggests that structural alerts associated with organ risk are a characteristic shared by most of the analyzed bioactive metabolites, rather than a feature unique to a subset. In contrast, nuclear receptor signaling and stress response pathways showed low toxicity predictions across the entire library. Isolated positive signals were restricted to individual compounds rather than cluster blocks or fungal origin. Overall, these results suggest that although the library exhibits significant predicted risk at the organ level, it does not demonstrate a generalized tendency toward endocrine disruption or genotoxic stress pathway activation.

Regarding xenobiotic metabolism, the evaluated cytochrome P450 (CYP) isoforms showed scattered positive signals distributed sporadically among individual compounds rather than concentrated in specific clusters. This distribution is consistent with the characteristic substrate specificity of each CYP isoform when tested against a chemically diverse library and does not necessarily reflect a systematic metabolic risk for the library as a whole.

### 3.2. Network pharmacology guided prioritization of autophagy-associated targets

After removing duplicate entries and integrating target predictions from PharmMapper and SwissTargetPrediction, 217 putative targets of fungal-derived compounds were identified. Additionally, 3,402 tauopathy-associated targets were obtained from DisGeNet, GeneCards, and STRING. Meanwhile, 8,847 autophagy-related targets were obtained from GeneCards and String. Intersecting these three datasets identified 91 overlapping targets of fungal-derived compounds, tauopathies, and autophagy as potential treatment targets for these neurodegenerative diseases (Figure 3A).

**Figure 3.**
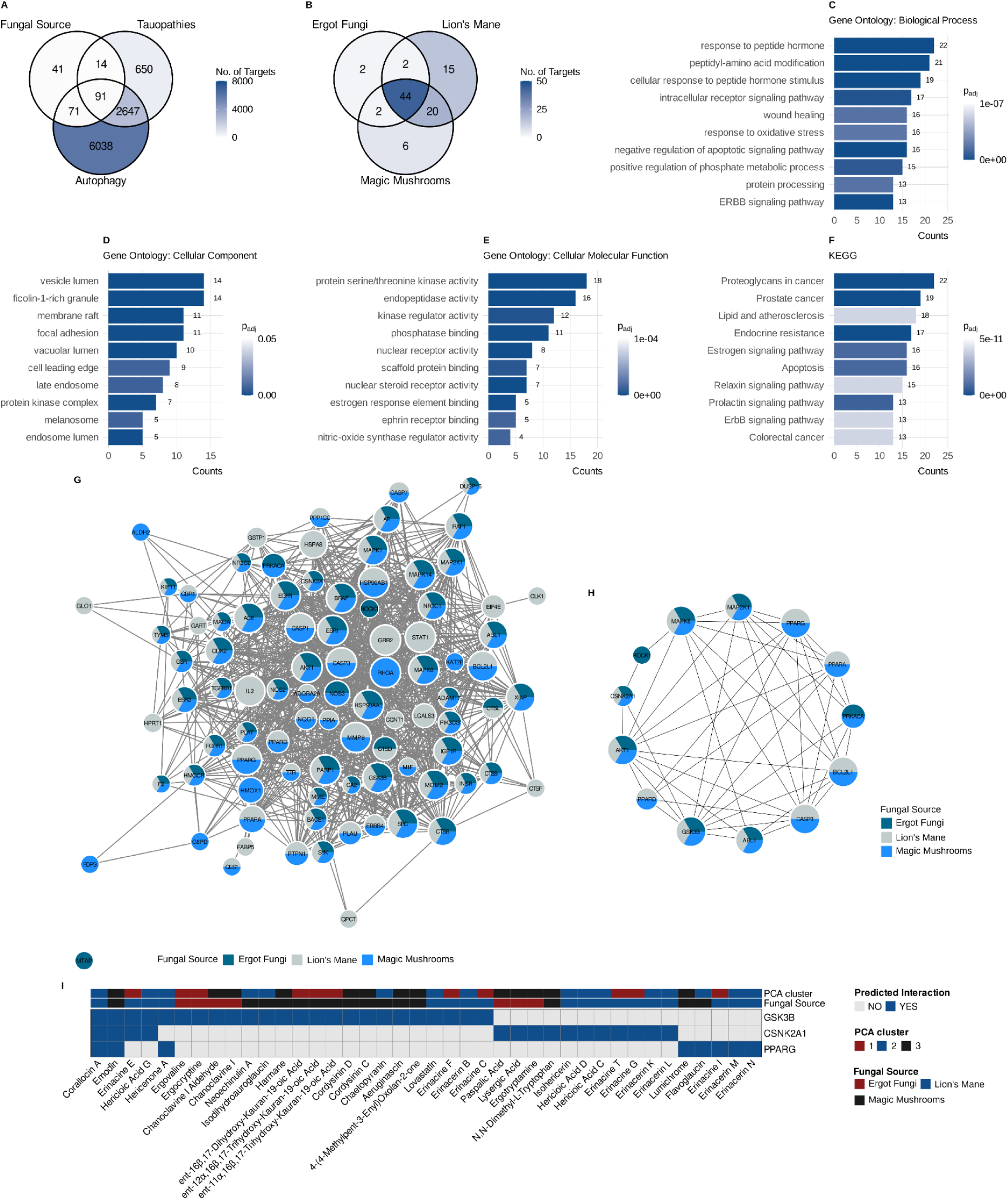
Identification and functional characterisation of molecular targets shared by neurobioactive fungal-derived compounds, tauopathies, and autophagy. (A) A Venn diagram showing the intersection between molecular targets associated with fungus-derived compounds, tauopathies and autophagy. This identifies the 91 shared targets that are used for subsequent analyses. (B) A Venn diagram showing the overlap of molecular targets amongst the three groups of fungi evaluated (Ergot Fungi, Lion’s Mane and Magic Mushrooms). (C–F) Functional enrichment analysis of the 91 shared targets using Gene Ontology for the terms’Biological Process’ (C),’Cellular Component’ (D) and’Molecular Function’ (E), as well as pathway enrichment analysis using KEGG (F). The length of the bars represents the number of genes associated with each term, while the colour intensity indicates the adjusted p-value (p_adj_). (G) The protein-protein interaction (PPI) network was constructed from the 91 shared targets. The size of the nodes is proportional to their degree of connectivity, and the coloured sectors indicate the fungal group to which each molecular target belongs. (H) A subnetwork comprising targets belonging to the Gene Ontology term’Regulation of autophagy’, supplemented with the PPARA, PPARD and PPARG isoforms due to their recognised role in transcriptional autophagy regulation. (I) Heat map showing predicted interactions between selected compounds and molecular targets prioritised for molecular docking and molecular dynamics studies. The annotations on the upside indicate the chemical cluster assigned to each compound and its fungal group of origin, while the colour indicates whether a predicted interaction is present or absent.

Of the 91 overlapping targets, 44 were shared by the three fungal sources (Figure 3B). This substantial overlap suggests that structurally diverse fungal metabolites may converge pharmacologically on a common set of molecular targets associated with autophagy and tauopathies.

Gene Ontology (GO) and KEGG enrichment analyses were performed to explore the biological functions of the 91 overlapping targets. GO enrichment analysis identified 357 significantly associated biological processes, 19 cellular components, and 47 molecular functions. KEGG analysis further revealed 159 significantly enriched pathways.

The top five terms for biological processes, ranked by adjusted p-value, were primarily enriched for response to peptide hormone (GO:0043434); peptidyl-amino acid modification (GO:0018193); cellular response to peptide hormone stimulus (GO:0071375); negative regulation of the apoptotic signalling pathway (GO:2001234); and the ERBB signalling pathway (GO:0038127) (Figure 3C). The top five terms for cellular components were ficolin-1-rich granule (GO:0101002), vesicle lumen (GO:0031983), vacuolar lumen (GO:0005775), membrane raft (GO:0045121), and endosome lumen (GO:0031904) (Figure 3D). The top five terms for molecular functions were: protein serine/threonine kinase activity (GO:0004674); nuclear steroid receptor activity (GO:0003707); endopeptidase activity (GO:0004175); nuclear receptor activity (GO:0004879); and estrogen response element binding (GO:0034056) (Figure 3E).

The top enriched KEGG pathways were mainly related to signal transduction, endocrine regulation, apoptosis, and cancer-associated pathways (Figure 3F). Notably, several pathways with well-established roles in autophagy regulation and neuronal survival were significantly enriched, including PI3K-Akt, MAPK, FoxO, ErbB, insulin, neurotrophin, Ras, and TNF signalling pathways. Overall, the enrichment analyses indicated that the overlapping targets participate predominantly in signalling pathways involved in cell survival, apoptosis, endocrine regulation, and neuronal function.

The PPI network constructed from the 91 overlapping targets, comprised 91 nodes and 1,020 edges (Figure 3G). To narrow this network down to the proteins that are most directly implicated in autophagy regulation, ten proteins that are annotated to the Gene Ontology term regulation of autophagy (GO:0010506) were identified: AKT1, CASP3, GSK3B, BCL2L1, MAPK8, ABL1, MAP2K1, PRKACA, CSNK2A1 and ROCK1. Given the well-documented role of peroxisome proliferator-activated receptors (PPARs) as transcriptional regulators of autophagy, the three PPAR isoforms present in the network were also included, resulting in a final subset of 13 targets for further characterization (Figure 3H).

Degree, betweenness, and closeness centrality were calculated for all 91 nodes of the PPI network, and the values corresponding to the 13 prioritized targets are summarized in Table 1. Within the full network, AKT1 ranked as the node with the highest connectivity among these 13 targets (degree = 72; betweenness = 899.99), followed by CASP3 (degree = 60, betweenness = 324.14) and, with equal degree, PPARG and GSK3B (degree = 49). PPARG and PPARA displayed disproportionately high betweenness centrality of 237.70 and 223.38, respectively; suggesting that these nuclear receptors may act as bridging nodes, connecting otherwise distinct regions of the broader network. In contrast, ROCK1 showed the lowest connectivity among the prioritized targets (degree = 8; betweenness = 1.25). Most of these autophagy and PPAR-related targets ranked among the most central nodes of the entire 91-target network, indicating convergence between the GO-based functional selection and the network’s intrinsic topological hierarchy.

**Table 1.** Topological properties of autophagy-modulating targets identified by network pharmacology analysis of neurobioactive fungal-derived compounds.

| Protein | Target name | Degree | Betweenness | Closeness |
| --- | --- | --- | --- | --- |
| AKT1 | RAC-alpha serine/threonine-protein kinase | 72 | 899.99 | 0.46 |
| CASP3 | Caspase-3 subunit p12 | 60 | 324.14 | 0.43 |
| PPARG | Peroxisome proliferator-activated receptor gamma | 49 | 237.70 | 0.41 |
| GSK3B | Glycogen synthase kinase-3 beta | 49 | 187.66 | 0.41 |
| BCL2L1 | Bcl-2-like protein 1 | 43 | 140.12 | 0.39 |
| MAPK8 | Mitogen-activated protein kinase 8 | 36 | 103.39 | 0.38 |
| ABL1 | Tyrosine-protein kinase ABL1 | 29 | 26.55 | 0.37 |
| MAP2K1 | Dual specificity mitogen-activated protein kinase kinase 1 | 29 | 16.58 | 0.37 |
| PPARA | Peroxisome proliferator-activated receptor alpha | 27 | 223.38 | 0.37 |
| PRKACA | cAMP-dependent protein kinase catalytic subunit alpha | 22 | 13.88 | 0.35 |
| PPARD | Peroxisome proliferator-activated receptor delta | 17 | 18.36 | 0.35 |
| CSNK2A1 | Casein kinase II subunit alpha | 15 | 14.25 | 0.35 |
| ROCK1 | Rho-associated protein kinase 1 | 8 | 1.25 | 0.33 |

Of the 13 prioritised targets, three were selected for molecular docking and molecular dynamics assays due to their mechanistic relevance to tau pathology and autophagy regulation: PPARG, GSK3B and CSNK2A1. PPARG was chosen as the PPAR isoform with the highest degree within the network (degree = 49), given its well-established role as a transcriptional regulator of autophagy-related genes. GSK3B was selected as a principal tau kinase and recognized autophagy modulator; inhibition of this enzyme is therefore expected to have a dual therapeutic effect by reducing pathological tau phosphorylation and promoting the clearance of tau aggregates via autophagy. CSNK2A1 was selected due to its well-documented role as a tau-associated kinase in tauopathies. Although its function as a negative regulator of autophagy initiation has been characterised in other pathological contexts, its involvement in autophagy regulation within tauopathies specifically remains largely unexplored. This makes it a novel target in the context of the present study.

Based on this selection, a total of 40 compounds with predicted interactions against PPARG, GSK3B, and/or CSNK2A1 were retained for molecular docking and molecular dynamics assays (Figure 3I). These compounds were distributed as follows: 8 compounds predicted to interact with PPARG, 24 with GSK3B, and 15 with CSNK2A1, with some compounds showing predicted interactions with more than one of the three targets.

### 3.3. Docking-based prioritization of fungal-derived compounds against autophagy-associated targets

The docking scores for these 40 compounds against their predicted targets are summarized in Table 2. Emodin, a Magic Mushroom-derived compound with previously characterised activity against all three targets, was included as an internal reference compound. This compound yielded binding affinities of-8.28,-9.57 and-9.64 kcal/mol against PPARG, GSK3B and CSNK2A1, respectively.

**Table 2.**
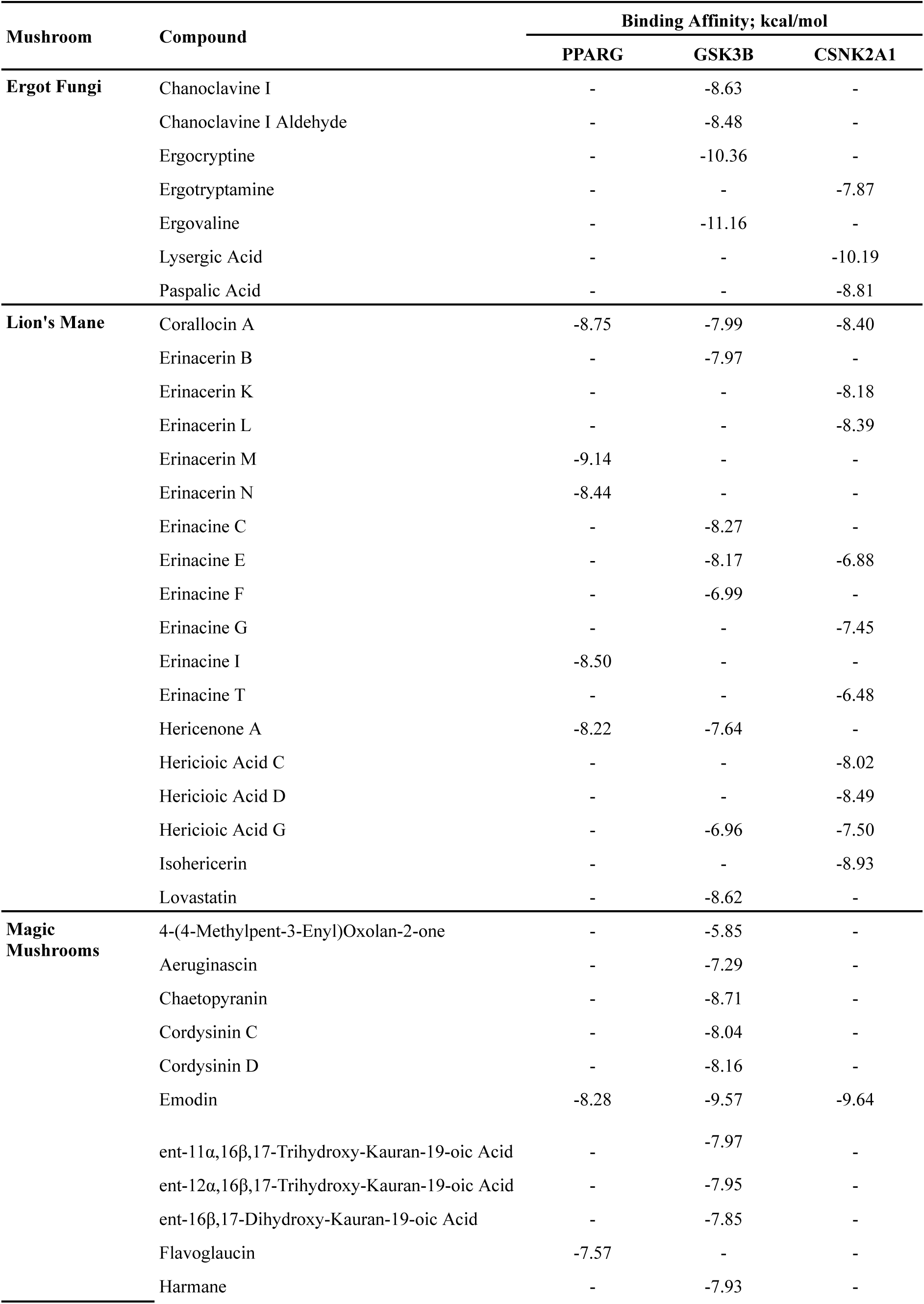
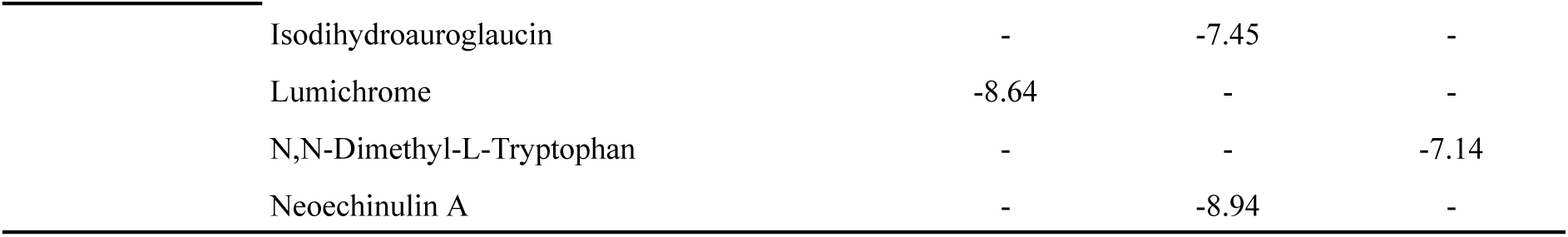
Binding affinities (kcal/mol) of neurobioactive fungal-derived compounds against PPARG, GSK3B, and CSNK2A1, corresponding to the best-ranked docking pose predicted by AutoDock Vina for each compound–target pair. . More negative values indicate stronger predicted binding.

| Mushroom | Compound | Binding Affinity; kcal/mol |  |  |
| --- | --- | --- | --- | --- |
|  |  | PPARG | GSK3B | CSNK2A1 |
| **Ergot Fungi** | Chanoclavine I | - | -8.63 | - |
|  | Chanoclavine I Aldehyde | - | -8.48 | - |
|  | Ergocryptine | - | -10.36 | - |
|  | Ergotryptamine | - | - | -7.87 |
|  | Ergovaline | - | -11.16 | - |
|  | Lysergic Acid | - | - | -10.19 |
|  | Paspalic Acid | - | - | -8.81 |
| **Lion's Mane** | Corallocin A | -8.75 | -7.99 | -8.40 |
|  | Erinacerin B | - | -7.97 | - |
|  | Erinacerin K | - | - | -8.18 |
|  | Erinacerin L | - | - | -8.39 |
|  | Erinacerin M | -9.14 | - | - |
|  | Erinacerin N | -8.44 | - | - |
|  | Erinacine C | - | -8.27 | - |
|  | Erinacine E | - | -8.17 | -6.88 |
|  | Erinacine F | - | -6.99 | - |
|  | Erinacine G | - | - | -7.45 |
|  | Erinacine I | -8.50 | - | - |
|  | Erinacine T | - | - | -6.48 |
|  | Hericenone A | -8.22 | -7.64 | - |
|  | Hericioic Acid C | - | - | -8.02 |
|  | Hericioic Acid D | - | - | -8.49 |
|  | Hericioic Acid G | - | -6.96 | -7.50 |
|  | Isohericerin | - | - | -8.93 |
|  | Lovastatin | - | -8.62 | - |
| **Magic Mushrooms** | 4-(4-Methylpent-3-Enyl)Oxolan-2-one | - | -5.85 | - |
|  | Aeruginascin | - | -7.29 | - |
|  | Chaetopyranin | - | -8.71 | - |
|  | Cordysinin C | - | -8.04 | - |
|  | Cordysinin D | - | -8.16 | - |
|  | Emodin | -8.28 | -9.57 | -9.64 |
| | ent-11α,16β,17-Trihydroxy-Kauran-19-oic Acid | - | -7.97 | - |
| | ent-12α,16β,17-Trihydroxy-Kauran-19-oic Acid | - | -7.95 | - |
| | ent-16β,17-Dihydroxy-Kauran-19-oic Acid | - | -7.85 | - |
|  | Flavoglaucin | -7.57 | - | - |
|  | Harmane | - | -7.93 | - |

|  |  |  |  |
| --- | --- | --- | --- |
| Isodihydroauroglaucin | - | -7.45 | - |
| Lumichrome | -8.64 | - | - |
| N,N-Dimethyl-L-Tryptophan | - | - | -7.14 |
| Neoechinulin A | - | -8.94 | - |

For PPARG, the strongest predicted affinities were observed for Erinacerin M (-9.14 kcal/mol), Corallocin A (-8.75 kcal/mol), and Lumichrome (-8.64 kcal/mol), followed by Erinacine I (-8.50 kcal/mol) and Erinacerin N (-8.44 kcal/mol). For GSK3B, Ergovaline (-11.16 kcal/mol) and Ergocryptine (-10.36 kcal/mol) produced the two strongest affinities across the entire dataset, followed by Neoechinulin A (-8.94 kcal/mol), Chaetopyranin (-8.71 kcal/mol), Chanoclavine I (-8.63 kcal/mol), and Lovastatin (-8.62 kcal/mol). For CSNK2A1, Lysergic Acid showed the strongest predicted affinity (-10.19 kcal/mol), followed by Isohericerin (-8.93 kcal/mol), Paspalic Acid (-8.81 kcal/mol), Hericioic Acid D (-8.49 kcal/mol), Corallocin A (-8.40 kcal/mol), and Erinacerin L (-8.39 kcal/mol).

These docking results were used to prioritise compounds for subsequent molecular dynamics (MD) simulations in descending order of predicted binding affinity for each target, beginning with the top-ranked hit. For each of the three targets, Emodin was simulated as an independent protein-ligand complex, regardless of its ranking within that target-specific set, allowing its results to serve as an internal benchmark for direct comparison against a ligand of known activity.

### 3.4. Molecular dynamics simulations of prioritized fungal-derived ligand–target complexes

For each of the three targets, the top-ranked docking solutions were used to build the corresponding receptor-ligand complexes, which were subjected to 300 ns molecular dynamics (MD) simulations. For each complex, we report the representative conformation of the most populous trajectory cluster, together with its corresponding hydrogen bonds and interaction distances; trajectory stability, assessed through pairwise RMSD heat maps; and the conformational space sampled during the simulation, examined by projecting the trajectory onto the first two principal components (Figures 4-6).

**Figure 4.**
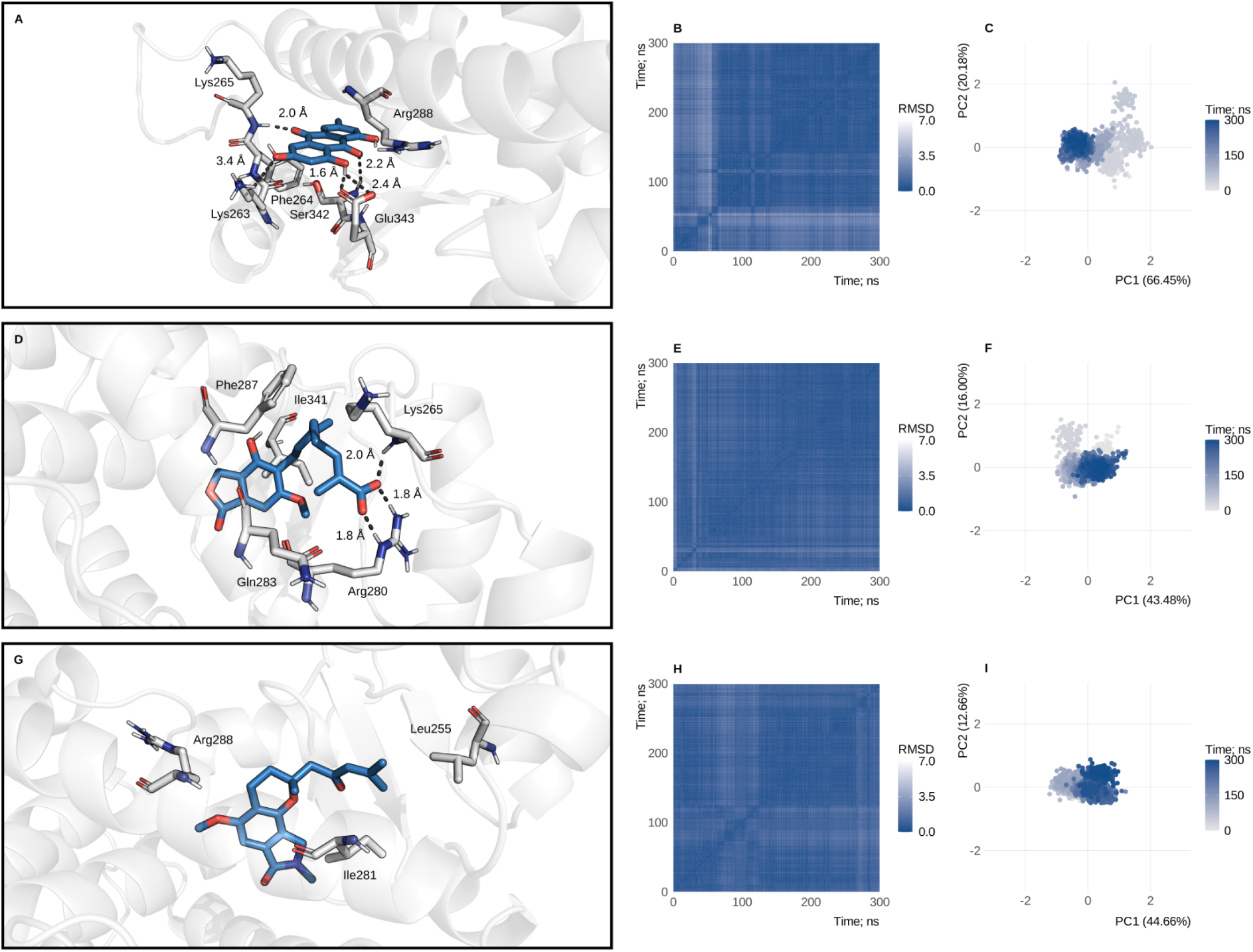
Validation of the predicted protein–ligand interactions for PPARG via molecular dynamics. (A–C): Results corresponding to the complex formed between Emodin and PPARG. (A) A representative conformation of the most populous cluster identified during the molecular dynamics simulation is shown, along with the main hydrogen bonds formed between the ligand and the residues at the binding site, and the interaction distances (Å). (B) Heat map of the root mean square deviation (RMSD) calculated from the molecular dynamics trajectory, where colour intensity represents the structural deviation between all pairs of sampled conformations. (C) Projection of the trajectory onto the first two principal components (PCA), where each coloured point represents an individual conformation and the percentages on the axes correspond to the fraction of total variance explained by each principal component. (D–F) Equivalent analyses for the PPARG–Corallocin A complex. (G–I) Equivalent analyses for the PPARG–Erinacerin M complex.

**Figure 5.**
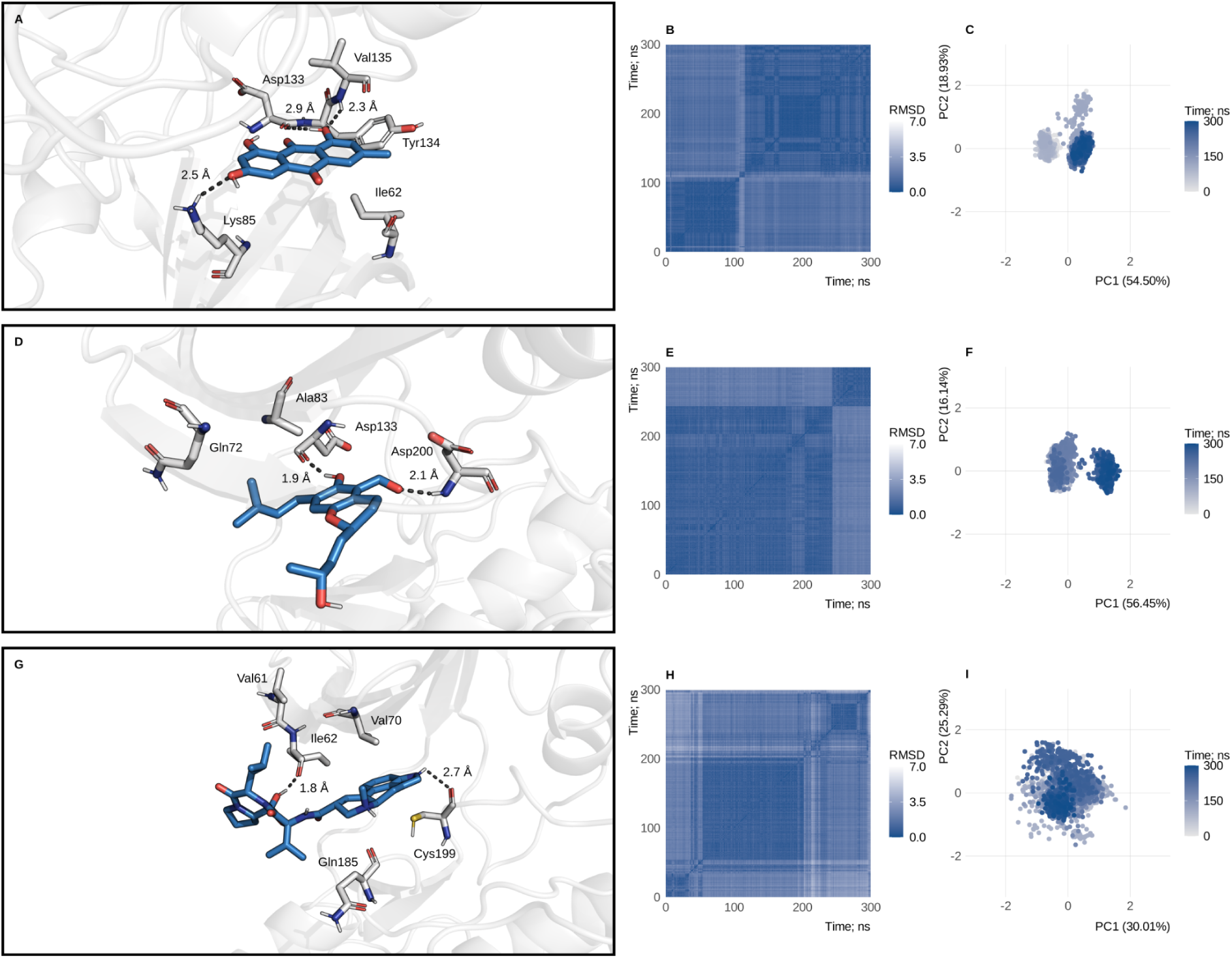
Validation of the predicted protein–ligand interactions for GSK3B via molecular dynamics. (A–C): Results corresponding to the complex formed between Emodin and GSK3B. (A) A representative conformation of the most populous cluster identified during the molecular dynamics simulation is shown, along with the main hydrogen bonds formed between the ligand and the residues at the binding site, and the interaction distances (Å). (B) Heat map of the root mean square deviation (RMSD) calculated from the molecular dynamics trajectory, where colour intensity represents the structural deviation between all pairs of sampled conformations. (C) Projection of the trajectory onto the first two principal components (PCA), where each coloured point represents an individual conformation and the percentages on the axes correspond to the fraction of total variance explained by each principal component. (D–F) Equivalent analyses for the GSK3B–Chaetopyranin complex. (G–I) Equivalent analyses for the GSK3B–Ergocryptine complex.

**Figure 6.**
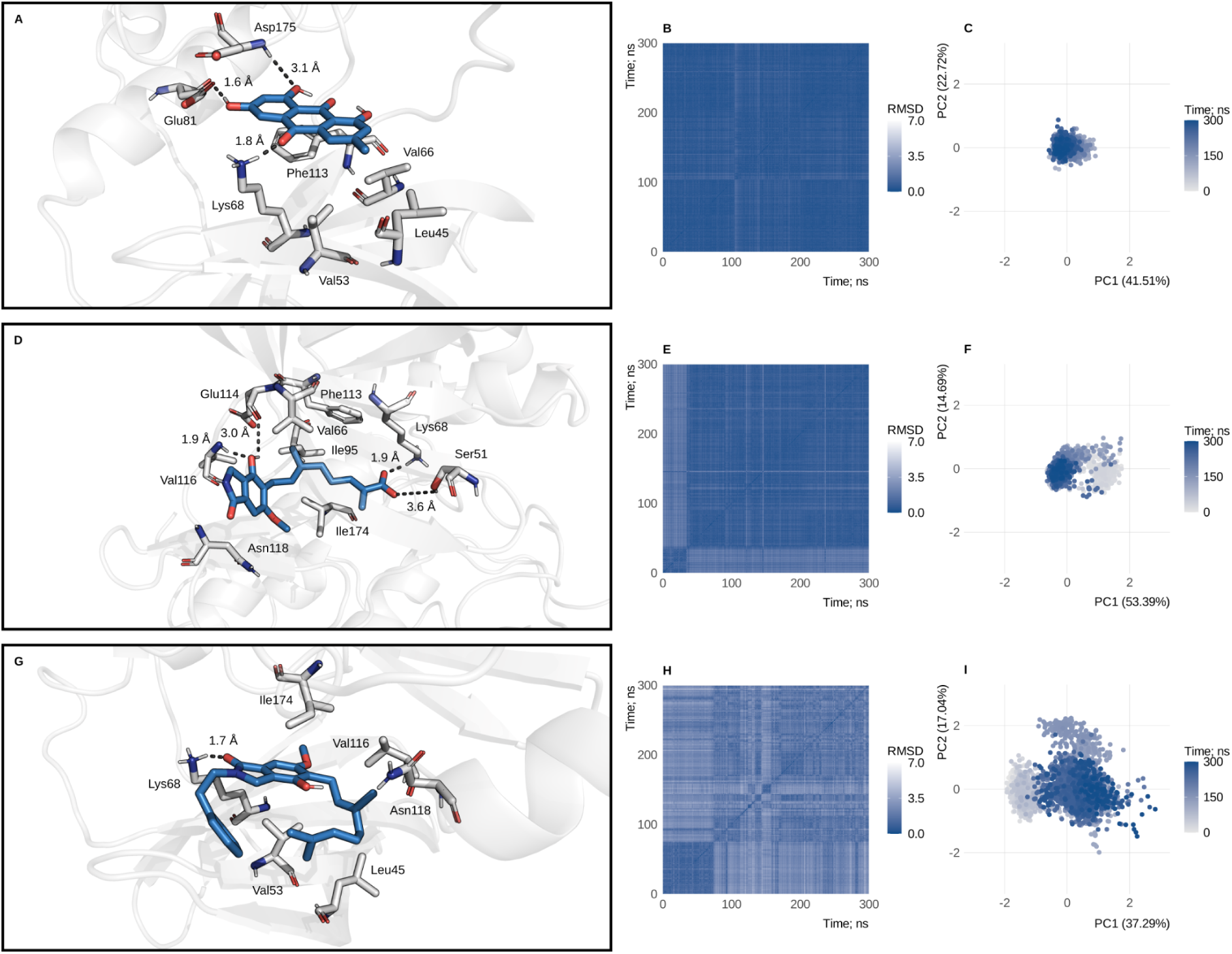
Validation of the predicted protein–ligand interactions for CSNK2A1 via molecular dynamics. (A–C): Results corresponding to the complex formed between Emodin and CSNK2A1. (A) A representative conformation of the most populous cluster identified during the molecular dynamics simulation is shown, along with the main hydrogen bonds formed between the ligand and the residues at the binding site, and the interaction distances (Å). (B) Heat map of the root mean square deviation (RMSD) calculated from the molecular dynamics trajectory, where colour intensity represents the structural deviation between all pairs of sampled conformations. (C) Projection of the trajectory onto the first two principal components (PCA), where each coloured point represents an individual conformation and the percentages on the axes correspond to the fraction of total variance explained by each principal component. (D–F) Equivalent analyses for the CSNK2A1–Hericioic Acid D complex. (G–I) Equivalent analyses for the CSNK2A1–Isohericerin complex.

#### 3.4.1. Stability of PPARG-ligand complexes during molecular dynamics simulations

To validate the predicted docking interactions for the top-ranked PPARG hits, MD simulations were performed on the complexes formed between PPARG and Corallocin A, Erinacerin M, and the internal reference compound Emodin (Figure 4).

In the PPARG-Emodin complex, the representative conformation of the most populous cluster was found to form a hydrogen bond with Ser342 and two hydrogen bonds with Glu343. It also formed a weak hydrogen bond with Lys263 and a hydrogen bond with Lys265. Meanwhile, Phe264 and Arg288 were found to engage with the ligand through hydrophobic interactions (Figure 4A). The RMSD heat map revealed a distinct region of increased structural deviation around the 50-60 ns interval, with comparatively lower and more consistent values throughout the remaining 300 ns of the trajectory (Figure 4B). Projecting onto the first two principal components, which accounted for 66.45% (PC1) and 20.18% (PC2) of the total variance respectively, revealed that conformations sampled during the first half of the trajectory were distributed towards positive PC1 values. In contrast, conformations sampled during the second half occupied a more compact region towards negative PC1 values (Figure 4C).

In the PPARG-Corallocin A complex, the representative conformation of the most populous cluster showed two hydrogen bonds with Arg280 and a hydrogen bond with Lys265, while Phe287, Ile341 and Gln283 engaged the ligand through hydrophobic interactions (Figure 4D). The RMSD heat map showed uniformly low values across the entire 300 ns trajectory, without discrete regions of increased deviation (Figure 4E). Projection onto the first two principal components, which accounted for 43.48% (PC1) and 16.00% (PC2) of the total variance, showed conformations from across the trajectory distributed within a single, comparatively compact region, with a subset of early conformations dispersed toward positive PC2 values (Figure 4F).

In the PPARG-Erinacerin M complex, the representative conformation of the largest cluster did not exhibit hydrogen bonding; rather, Arg288 interacted with the ligand via hydrophobic contact through its aliphatic side chain, and Leu255 and Ile281 also interacted with the ligand through hydrophobic interactions (Figure 4G). The RMSD heat map showed uniformly low values throughout the 300 ns trajectory, comparable to the pattern observed for the Corallocin A complex (see Figure 4H). Projecting onto the first two principal components, which together accounted for 44.66% (PC1) and 12.66% (PC2) of the total variance, revealed that conformations from across the entire trajectory were distributed within a single compact cluster (Figure 4I).

#### 3.4.2. Stability of GSK3B-ligand complexes during molecular dynamics simulations

To validate the predicted docking interactions of the top-ranked GSK3B hits, MD simulations were performed on complexes formed between GSK3B, the internal reference compound Emodin, and the top-ranked docking hits, selected in descending order of docking affinity. However, Ergovaline was excluded from further analysis after an initial MD simulation showed the ligand dissociating from the binding pocket, despite showing the highest predicted affinity for GSK3B (Table 2). Chaetopyranin and Ergocryptine, which remained stably bound throughout the simulation, were therefore selected for the analyses presented in this section (Figure 5).

In the GSK3B-Emodin complex, the representative conformation of the largest cluster was found to form three hydrogen bonds: one with Asp133, one with Val135 and one with Lys85. Meanwhile, Ile62 and Tyr134 formed hydrophobic interactions with the ligand (Figure 5A). The RMSD heat map revealed two distinct regions of low internal deviation, divided by a transition at around 110 ns. This indicates a shift between two primary conformational states throughout the trajectory (Figure 5B). Projection onto the first two principal components, which accounted for 54.50% (PC1) and 18.93% (PC2) of the total variance, revealed that conformations sampled before and after this transition occupied two adjacent yet distinguishable regions along PC1 (Figure 5C).

In the GSK3B-Chaetopyranin complex, the representative conformation of the most populous cluster formed hydrogen bonds with Asp133 and Asp200, while Gln72 and Ala83 interacted with the ligand through hydrophobic interactions (Figure 5D). The RMSD heat map showed a comparatively minor transition at around 250 ns, with uniform low deviation otherwise across the trajectory (Figure 5E). Projecting onto the first two principal components, which accounted for 56.45% (PC1) and 16.14% (PC2) of the total variance respectively, revealed two adjacent, partially overlapping regions along PC1, which was consistent with the RMSD pattern (see Figure 5F).

In the GSK3B-Ergocryptine complex, the representative conformation of the most populous cluster formed hydrogen bonds with Ile62 and Cys199, while Val61, Val70, and Gln185 interacted with the ligand through hydrophobic forces (Figure 5G). The RMSD heat map showed uniformly low values across most of the 300 ns trajectory, with a discrete, transient region of increased deviation around 240-280 ns (Figure 5H). Projecting onto the first two principal components, which accounted for 30.01% (PC1) and 25.29% (PC2) of the total variance respectively, revealed that the conformations were distributed within a single, broad, continuous region, with no clear separation between the early and late time points (Figure 5I).

#### 3.4.3. Stability of CSNK2A1-ligand complexes during molecular dynamics simulations

To validate the predicted docking interactions of the top-ranked CSNK2A1 hits, MD simulations were performed on complexes formed between CSNK2A1, the internal reference compound Emodin, and the top-ranked docking hits, selected in descending order of docking affinity. However, Lysergic Acid was excluded from further analysis after an initial MD simulation showed the ligand dissociating from the binding pocket, despite showing the highest predicted affinity for CSNK2A1 (Table 2). Hericioic Acid D and Isohericerin, which remained stably bound throughout the simulation, were therefore selected for the analyses presented in this section (Figure 6).

In the CSNK2A1-Emodin complex, the representative conformation of the largest cluster was found to form three hydrogen bonds: one with Lys68, one with Glu81 and a weaker hydrogen bond with Asp175. Meanwhile, Leu45, Val53, Val66 and Phe113 formed hydrophobic interactions with the ligand (Figure 6A). The RMSD heat map showed uniformly low values across the entire 300 ns trajectory without any regions of increased deviation (Figure 6B). Projecting onto the first two principal components, which accounted for 41.51% (PC1) and 22.72% (PC2) of the total variance respectively, revealed that conformations from across the trajectory were distributed within a single, compact region (Figure 6C).

In the CSNK2A1-Hericioic Acid D complex, the representative conformation of the most populous cluster was found to form a hydrogen bond with Lys68 and with Val116. It also formed a weak hydrogen bond with Ser51 and with Glu114. Meanwhile, Val66, Ile95, Phe113, Asn118 and Ile174 were found to engage with the ligand through hydrophobic interactions (Figure 6D). The RMSD heat map showed a transition around 40 ns, with comparatively uniform, low deviation before and after this point (Figure 6E). Projecting onto the first two principal components, which accounted for 53.39% (PC1) and 14.69% (PC2) of the total variance, revealed that the conformations sampled during the initial quarter of the trajectory were distributed towards positive PC1 values. Meanwhile, the remaining conformations occupied a more compact, adjacent region (Figure 6F).

In the CSNK2A1-Isohericerin complex, the representative conformation of the most populous cluster was found to form a hydrogen bond with Lys68. Meanwhile, Leu45, Val53, Val116, Asn118 and Ile174 were found to engage with the ligand through hydrophobic interactions (Figure 6G). The RMSD heat map showed a discrete region of increased deviation around 70–100 ns, with comparatively lower and more uniform values across the remainder of the trajectory (Figure 6H). Projection onto the first two principal components, which accounted for 37.29% (PC1) and 17.04% (PC2) of the total variance, showed conformations sampled during approximately the first quarter of the trajectory distributed toward negative PC1 and positive PC2 values, while the remaining conformations occupied a broader, more central region (Figure 6I).

### 3.5. MM/GBSA binding free energy calculations of prioritized ligand–target complexes

MM/GBSA calculations were performed on the MD trajectories of the ligand-protein complexes simulated for the three targets, providing a more rigorous, ensemble-based estimate of binding free energy to complement the rankings obtained from docking (Table 3). Emodin, included as a reference compound, consistently displayed the weakest binding free energies across all three targets (-31.53 ± 5.23,-23.43 ± 4.25, and-24.23 ± 2.77 kcal/mol for PPARG, GSK3B, and CSNK2A1, respectively), indicating that the prioritized fungal metabolites generally outperformed it in terms of binding stability. A notable exception was observed for PPARG, where Erinacerin M showed a binding free energy statistically comparable to emodin.

**Table 3.** MM/GBSA binding free energy calculations for the protein–ligand complexes. Values are expressed as mean ± standard deviation (kcal/mol). ΔE_vdW_, van der Waals energy contribution; ΔE_ele_, electrostatic energy contribution; ΔG_bind_, total binding free energy estimated by the MM/GBSA method. Values are presented as mean ± standard deviation calculated from the MD trajectories.

| Target | Ligand | $\Delta E_{\text{vdW}}$; kcal/mol | $\Delta E_{\text{ele}}$; kcal/mol | $\Delta G_{\text{bind}}$; kcal/mol |
| --- | --- | --- | --- | --- |
| **PPARG** | Emodin | $-35.40 \pm 3.73$ | $-54.46 \pm 21.94$ | $-31.53 \pm 5.23$ |
| | Corallocin A | $-45.32 \pm 3.34$ | $-61.76 \pm 10.63$ | $-54.26 \pm 4.51$ |
| | Erinacerin M | $-46.06 \pm 2.47$ | $-5.10 \pm 3.07$ | $-31.61 \pm 2.88$ |
| **GSK3B** | Emodin | $-30.97 \pm 3.78$ | $-22.05 \pm 7.24$ | $-23.43 \pm 4.25$ |
| | Chaetopyranin | $-40.35 \pm 3.23$ | $-24.30 \pm 6.76$ | $-36.01 \pm 4.82$ |
| | Ergocryptine | $-50.32 \pm 4.49$ | $15.48 \pm 11.13$ | $-35.79 \pm 4.34$ |
| **CSNK2A1** | Emodin | $-31.30 \pm 2.67$ | $-31.21 \pm 4.07$ | $-24.23 \pm 2.77$ |
| | Hericioic Acid D | $-42.52 \pm 3.36$ | $-111.23 \pm 19.69$ | $-42.39 \pm 4.07$ |
| | Isohericerin | $-41.17 \pm 4.56$ | $-2.22 \pm 5.07$ | $-28.81 \pm 5.87$ |

For PPARG, Corallocin A exhibited the most favourable binding free energy (ΔG_bind_ =-54.26 ± 4.51 kcal/mol), which was nearly twice as favourable as that of Emodin and Erinacerin M (-31.61 ± 2.88 kcal/mol). This enhanced affinity was driven by substantial contributions from van der Waals (-45.32 ± 3.34 kcal/mol) and electrostatic (-61.76 ± 10.63 kcal/mol) interactions. Erinacerin M demonstrated a van der Waals contribution (-46.06 ± 2.47 kcal/mol) comparable to that of Corallocin A, yet its significantly weaker electrostatic term (-5.10 ± 3.07 kcal/mol) resulted in an overall binding free energy that was similar to that of Emodin. This highlights the importance of electrostatic complementarity in differentiating high-affinity from moderate-affinity ligands for this target.

For GSK3B, both Chaetopyranin and Ergocryptine showed improved binding relative to emodin, with ΔG_bind_ values of-36.01 ± 4.82 and-35.79 ± 4.34 kcal/mol, respectively. These two ligands achieved comparable overall affinities through different energetic profiles: Chaetopyranin combined moderate van der Waals (-40.35 ± 3.23 kcal/mol) and electrostatic (-24.30 ± 6.76 kcal/mol) contributions, whereas Ergocryptine relied almost entirely on a strong van der Waals term (-50.32 ± 4.49 kcal/mol) to compensate for an unfavorable electrostatic contribution (15.48 ± 11.13 kcal/mol). This suggests that hydrophobic packing can offset unfavorable electrostatics to sustain overall binding stability in the GSK3B pocket.

For CSNK2A1, Hericioic Acid D showed the most favorable binding free energy (-42.39 ± 4.07 kcal/mol), outperforming Emodin and Isohericerin (-28.81 ± 5.87 kcal/mol). This result was dominated by an exceptionally strong electrostatic contribution (ΔE_ele_ =-111.23 ± 19.69 kcal/mol), the largest observed among all complexes analyzed, together with a favorable van der Waals term (-42.52 ± 3.36 kcal/mol). In contrast, Isohericerin, despite a comparable van der Waals contribution (-41.17 ± 4.56 kcal/mol), showed a negligible electrostatic term (-2.22 ± 5.07 kcal/mol), leading to a substantially weaker overall binding free energy.

The results show that van der Waals interactions are the main energetic driver of binding across all three targets, which is consistent with predominantly hydrophobic binding pockets. However, favourable electrostatic contributions further enhanced binding stability in the top-performing complexes for each target (PPARG-Corallocin A, GSK3B-Chaetopyranin and CSNK2A1-Hericioic Acid D), reinforcing their prioritisation over emodin and the remaining candidate ligands. Thus, these MM/GBSA results, calculated over the MD trajectories of each complex, provide energetic support for the ligand ranking obtained from docking and strengthen confidence in the selected fungal metabolites as promising candidates for further evaluation.

### 3.6. Complementary ligand-based QSAR modelling

To complement the docking, molecular dynamics, and MM/GBSA analyses, machine learning-based QSAR models were developed to predict the binding affinity (pKd or pKi, depending on the target) of fungal-derived compounds against PPARG, GSK3B, and CSNK2A1, using molecular descriptors derived from the optimized ligand structures as model inputs. Six regression algorithms were evaluated (Elastic Net, Random Forest, Support Vector Regression, Cubist, Gradient Boosting Machine, and Partial Least Squares), and performance was assessed based on training-set fit (R², MAE, RMSE), internal cross-validation (*Q^2^_CV_*), and external test-set performance (R², MAE, RMSE) (Table 4). For each target, the final model was not solely selected on the basis of the highest testing R², but rather by prioritising models that combined strong *Q^2^_CV_* values with consistent external test-set performance. This is because achieving this balance reduces the risk of selecting an overfitted model that only performs well on a particular training/test partition.

**Table 4.** Performance metrics of the machine learning-based QSAR models developed for PPARG, GSK3B, and CSNK2A1. Data are presented as coefficient of determination (R²), cross-validated coefficient of determination (*Q^2^_CV_*), mean absolute error (MAE), and root mean square error (RMSE). Models marked with an asterisk (*) were selected as the final predictive models for each target based on their external validation performance.

| Target | Model | Training | | | $Q_{cv}^2$ | Testing | | |
| --- | --- | --- | --- | --- | --- | --- | --- | --- |
| | | MAE | RMSE | $R^2$ | | MAE | RMSE | $R^2$ |
| PPARG | Elastic Net | 0.37 | 0.46 | 0.88 | 0.68 | 0.60 | 0.69 | 0.81 |
|  | Random Forest | 0.33 | 0.43 | 0.92 | 0.65 | 0.53 | 0.66 | 0.75 |
|  | Support Vector Regression* | 0.21 | 0.31 | 0.95 | 0.72 | 0.48 | 0.53 | 0.85 |
|  | Cubist | 0.34 | 0.41 | 0.91 | 0.55 | 0.53 | 0.67 | 0.72 |
|  | Gradient Boosting Machine | 0.25 | 0.30 | 0.95 | 0.62 | 0.57 | 0.65 | 0.74 |
|  | Partial Least Squares | 0.38 | 0.49 | 0.88 | 0.68 | 0.69 | 0.79 | 0.68 |
| GSK3B | Elastic Net | 0.39 | 0.53 | 0.64 | 0.44 | 0.43 | 0.59 | 0.51 |
|  | Random Forest | 0.16 | 0.23 | 0.95 | 0.54 | 0.36 | 0.53 | 0.62 |
|  | Support Vector Regression | 0.18 | 0.31 | 0.88 | 0.55 | 0.38 | 0.55 | 0.59 |
|  | Cubist* | 0.13 | 0.18 | 0.96 | 0.57 | 0.35 | 0.53 | 0.62 |
|  | Gradient Boosting Machine | 0.23 | 0.32 | 0.88 | 0.54 | 0.37 | 0.55 | 0.58 |
|  | Partial Least Squares | 0.51 | 0.65 | 0.45 | 0.22 | 0.48 | 0.64 | 0.44 |
| CSNK2A1 | Elastic Net | 0.57 | 0.75 | 0.62 | 0.43 | 0.62 | 0.91 | 0.53 |
|  | Random Forest | 0.25 | 0.34 | 0.94 | 0.58 | 0.55 | 0.75 | 0.66 |
|  | Support Vector Regression | 0.20 | 0.36 | 0.92 | 0.52 | 0.56 | 0.74 | 0.68 |
|  | Cubist | 0.23 | 0.32 | 0.93 | 0.53 | 0.55 | 0.82 | 0.63 |
|  | Gradient Boosting Machine* | 0.19 | 0.28 | 0.95 | 0.60 | 0.53 | 0.75 | 0.66 |
|  | Partial Least Squares | 0.61 | 0.79 | 0.57 | 0.41 | 0.60 | 0.88 | 0.60 |

Support Vector Regression (SVR) was selected as the final PPARG model, showing the highest *Q^2^_CV_* value (0.72) and the best external validation performance (R² = 0.85, MAE = 0.48, RMSE = 0.53) of the six tested algorithms. The agreement between the results of the internal cross-validation and the external testing, together with the relatively small difference between the training and testing performance (R² = 0.95 and R² = 0.85, respectively), suggests that SVR generalised well beyond the training set. This is in contrast to tree-based alternatives, such as Random Forest and Gradient Boosting Machine, which achieved comparably high training R² values, but lower *Q^2^_CV_* values and weaker testing performance.

For GSK3B, Cubist was selected as the final model, combining the highest *Q^2^_CV_* (0.57) with a testing R² of 0.62, matched only by Random Forest, while achieving the lowest testing MAE (0.35) among all algorithms evaluated. The consistency between cross-validated and external performance results in favour of Cubist as the most robust choice for this target, in contrast to models such as Elastic Net and Partial Least Squares, which exhibited lower *Q^2^_CV_* values and weaker testing metrics.

For CSNK2A1, Gradient Boosting Machine (GBM) was selected as the final model based on its markedly higher *Q^2^_CV_* (0.60) compared to all other algorithms, including Support Vector Regression (*Q^2^_CV_*=0.52), which showed a marginally higher testing R² (0.68 vs. 0.66) and lower testing RMSE (0.74 vs. 0.75) than GBM. As a lower *Q^2^_CV_* can indicate a model that is more sensitive to the specific training/test partition and therefore more susceptible to overfitting, the superior cross-validated performance of GBM, combined with testing metrics comparable to those of the most effective alternative, justified its selection as the more reliable and generalizable predictor for this target.

The predictive performance of the three final models was examined graphically (Figure 7). Scatter plots of experimental versus predicted binding affinities (Figures 7A, D and G) showed that the distribution of both training and external test compounds was close to the identity line for all three targets. This indicates that there was no systematic bias towards over-or underprediction and supports the adequacy of the SVR, Cubist and GBM models for PPARG, GSK3B and CSNK2A1, respectively.

**Figure 7.**
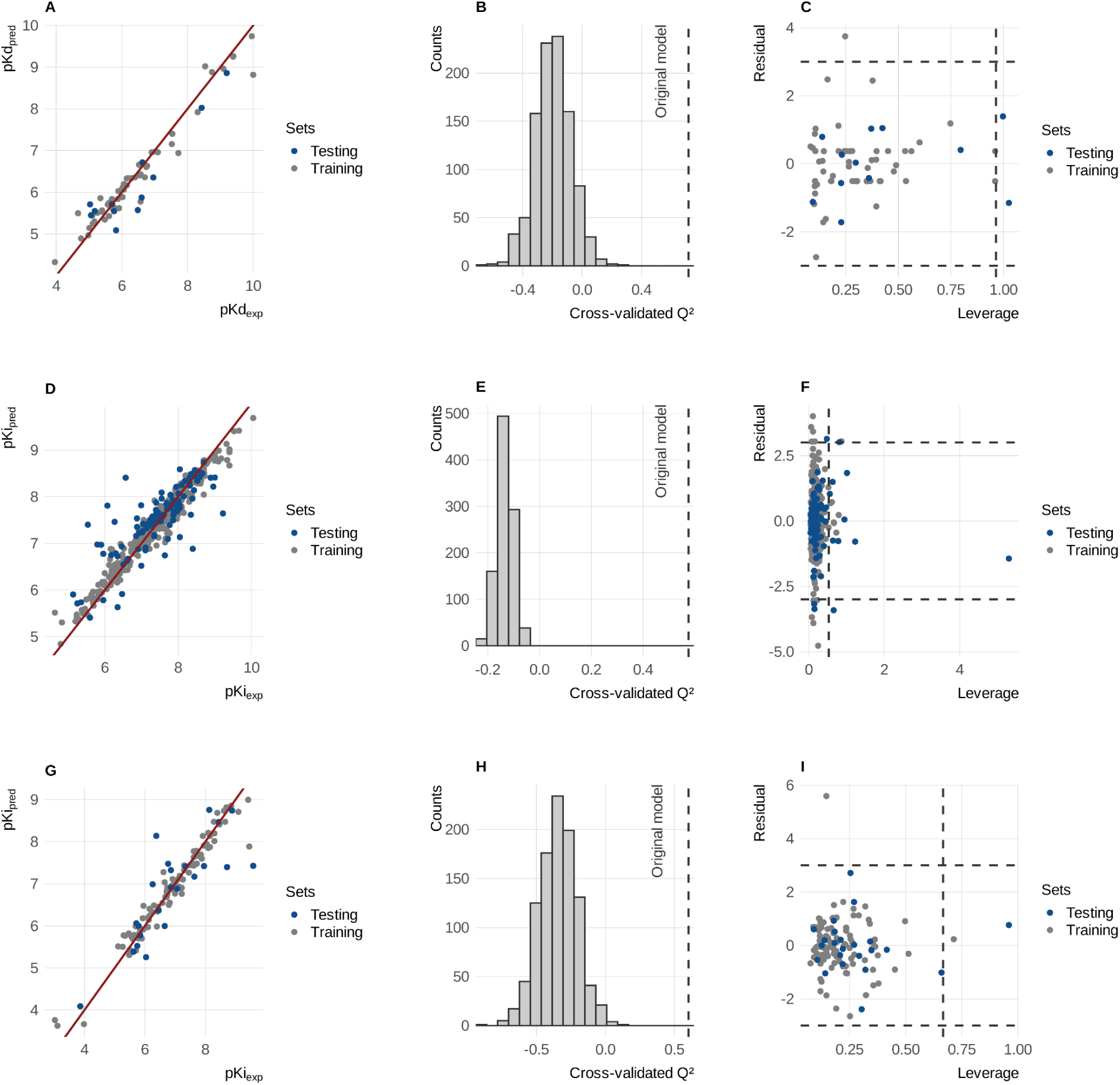
Validation of the final QSAR models developed for PPARG, GSK3B, and CSNK2A1. (A, D, G) Scatter plots comparing experimental and predicted binding affinities for the PPARG (pKd), GSK3B (pKi), and CSNK2A1 (pKi) models, respectively. Gray and blue points represent the training and external test sets, respectively, while the red line denotes the identity line (y = x). (B, E, H) Y-randomization results showing the distribution of cross-validated Q² values obtained from randomly permuted response variables. The dashed vertical line indicates the Q² value of the original model. (C, F, I) Williams plots used to assess the applicability domain of each QSAR model. The horizontal dashed lines indicate the standardized residual threshold (±3), whereas the vertical dashed line represents the warning leverage (h*).

The robustness of the models was also assessed using Y-randomisation, whereby the cross-validated Q² values of each original model were compared with the distribution of Q² values obtained by repeatedly permuting the response variable (Figures 7B, E and H). In all three cases, the Q² value of the original model fell well outside the range of the permuted distributions, which clustered around zero or below. This confirmed that the observed correlations between molecular descriptors and biological activity were not due to chance, and that each model captured a genuine structure-activity relationship.

Finally, the applicability domain of each model was evaluated using Williams plots (Figures 7C, F and I). For PPARG, only one training compound exceeded the ±3 residual threshold, while two external test compounds exceeded the warning leverage limit. One external test compound for GSK3B displayed a leverage value substantially higher than the rest of the dataset, identifying it as a structurally influential outlier. Predictions for this compound should be interpreted with caution when extrapolating to chemically dissimilar compounds. A second external test compound was found to marginally exceed the ±3 residual threshold. Similarly, one training compound for CSNK2A1 exceeded the residual threshold, indicating comparatively poor prediction despite falling within the leverage-based structural domain.Overall, the vast majority of training and test compounds across all three targets remained within the combined leverage/residual boundaries, supporting the internal consistency of the three QSAR models within the chemical space they were trained on. Whether the fungal-derived candidates of interest fall within this same space is addressed in the following section.

### 3.7. QSAR prediction of fungal-derived ligands

We subsequently applied the three QSAR models to estimate the binding affinities of a set of fungal-derived compounds against PPARG, GSK3B, and CSNK2A1, with the corresponding applicability domain (AD) classification reported in Table 5 to indicate the reliability of each individual prediction.

**Table 5.** QSAR predictions and applicability domain assessment for selected fungal-derived compounds targeting PPARG, GSK3B, and CSNK2A1. Predicted binding affinities are reported as pKd for PPARG and pKi for GSK3B and CSNK2A1. The applicability domain was assessed using the Williams plot. Predictions classified as *Inside* indicate compounds located within the structural applicability domain of the corresponding QSAR model, whereas *Outside* indicates compounds lying outside the model’s chemical space and whose predicted affinities should therefore be interpreted with caution.

| Compound | PPARG |  | GSK3B |  | CSNK2A1 |  |
| --- | --- | --- | --- | --- | --- | --- |
|  | pKd | Applicability Domain | pKi | Applicability Domain | pKi | Applicability Domain |
| Emodin | 5.85 | Inside | 6.90 | Outside | 6.38 | Inside |
| Corallocin A | 4.52 | Inside | - | - | - | - |
| Erinacerin M | 4.61 | Inside | - | - | - | - |
| Chaetopyranin | - | - | 6.53 | Outside | - | - |
| Ergocryptine | - | - | 4.67 | Outside | - | - |
| Hericioic Acid D | - | - | - | - | 6.09 | Outside |
| Isohericerin | - | - | - | - | 6.67 | Outside |

Emodin showed a moderate predicted affinity towards both PPARG (pKi = 5.85) and CSNK2A1 (pKi = 6.38). Both of these predictions fell within the structural applicability domain of their respective models, meaning they are considered reliable. However, its predicted affinity towards GSK3B (pKi = 6.90) fell outside the model’s applicability domain. Despite Emodin’s established bioactivity profile, this particular prediction should therefore be interpreted with caution.

Of the remaining fungal-derived compounds, Corallocin A and Erinacerin M were evaluated exclusively against PPARG. They yielded lower predicted affinities (pKi = 4.52 and 4.61, respectively) and were both located within the applicability domain. For GSK3B, Chaetopyranin and Ergocryptine returned predicted pKi values of 6.53 and 4.67, respectively. However, like emodin, both compounds were classified as lying outside the applicability domain, suggesting that this region of chemical space is generally underrepresented in the training set used to build the GSK3B model. A similar pattern was observed for CSNK2A1; Hericioic Acid D and Isohericerin exhibited the highest predicted affinities in the entire dataset (pKi = 6.09 and 6.67, respectively); however, both compounds fell outside the model’s applicability domain. This warrants cautious interpretation, despite their promising affinity values.

Overall, the inclusion of Emodin as a control allowed the predictions for the novel fungal ligands to be benchmarked against a compound with known activity. Corallocin A and Erinacerin M were the only candidates for which the QSAR prediction fell within the applicability domain of the PPARG model, providing an additional, independent line of support consistent with their docking, MD, and MM/GBSA results. For Chaetopyranin, Ergocryptine, Hericioic Acid D, and Isohericerin, the QSAR predictions fell outside the corresponding models’ applicability domains and are therefore not informative either for or against their prioritization; their selection as candidates continues to rest on the converging structural evidence from docking, molecular dynamics, and MM/GBSA described above.

### 3.8. Reanalysis of human hippocampal proteomic data

To provide independent, patient-derived support for the autophagy-tauopathy link identified through the above-described network-based analyses, we reanalyzed publicly available quantitative proteomic data from the hippocampus of AD patients. Since this dataset included proteomic profiling from multiple hippocampal subfields, differential expression analysis was performed for each region available. Significant enrichment of autophagy-and lysosome-related GO and KEGG terms was observed exclusively in the CA3 subfield; no significant enrichment of these terms was detected in the remaining hippocampal regions analysed. As noted in Methods, this region-by-region comparison was not corrected for the number of subfields tested, and the CA3-specific finding is therefore reported as an exploratory observation rather than a confirmed regional effect. Subsequent analyses therefore focused specifically on the CA3 region.

Within CA3, differentially expressed proteins (DEPs) were identified using two independent TMT-based quantitative proteomic experiments, a 10-plex and a 6-plex set. The 10-plex experiment yielded 532 DEPs exclusively, while the 6-plex experiment yielded 2,890 exclusively; 665 DEPs were common to both experiments (Figure 8A). Therefore, these 665 overlapping proteins were selected for downstream functional enrichment analysis.

**Figure 8.**
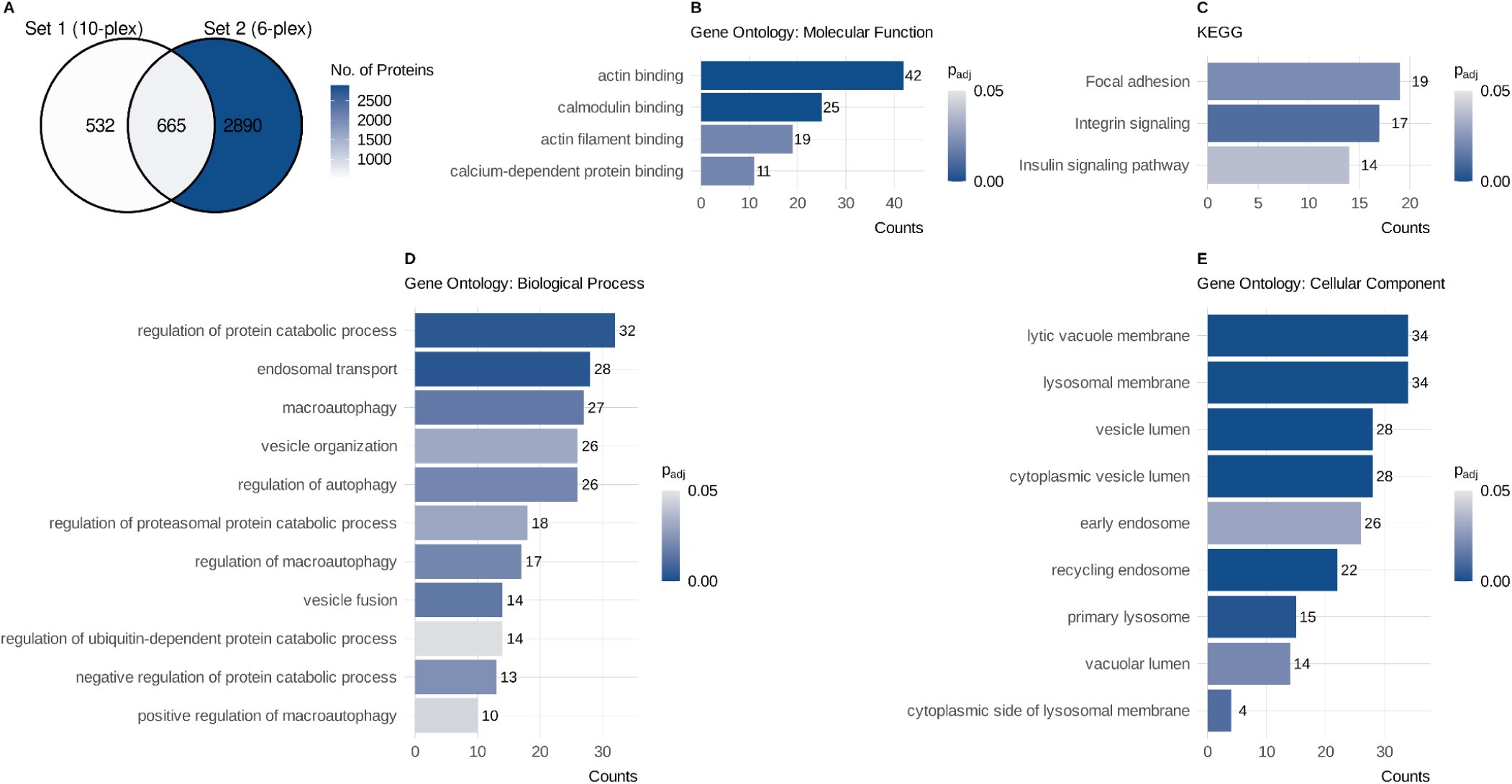
Proteomic evidence of autophagy dysregulation in the CA3 region of the hippocampus from patients with Alzheimer’s disease. (A) Overlap of differentially expressed proteins identified in the CA3 region using 10-plex and 6-plex TMT-based quantitative proteomics. (B–E) Functional enrichment analysis of the shared differentially expressed proteins showing significantly enriched Gene Ontology Molecular Function (B), KEGG pathways (C), Gene Ontology Biological Process (D), and Gene Ontology Cellular Component (E) categories. Bar length indicates the number of proteins assigned to each term, whereas color intensity represents the adjusted p-value (p_adj_).

GO and KEGG enrichment analyses of these 665 shared DEPs identified 74 significantly enriched biological processes, 61 cellular components, and 8 molecular functions, together with 8 significantly enriched KEGG pathways. Among these terms, a subset directly related to autophagy and lysosomal degradation was identified. Regarding molecular function, this subset included actin binding, calmodulin binding, actin filament binding, and calcium-dependent protein binding (Figure 8B). These functions are all linked to cytoskeletal dynamics and vesicular trafficking. Among the enriched KEGG pathways, those relevant to autophagy regulation included focal adhesion, integrin signaling, and the insulin signaling pathway (Figure 8C).

Within the biological process category, several enriched terms were directly related to autophagy. These terms included macroautophagy, regulation of autophagy, regulation of macroautophagy, and positive regulation of macroautophagy. Related processes included endosomal transport, vesicle organization, vesicle fusion, and regulation of the protein catabolic process, including its proteasomal and ubiquitin-dependent subcategories (Figure 8D). Similarly, several of the enriched cellular component terms corresponded to organelles central to the autophagic-lysosomal pathway. These organelles include the lysosomal membrane, lytic vacuole membrane, primary lysosome, vacuolar lumen, early and recycling endosomes, and vesicle/cytoplasmic vesicle lumen (Figure 8E). Taken together, these enrichment results indicate that autophagy-and lysosome-related processes are altered in the CA3 region of the hippocampus in patients with AD, consistent with the autophagy-tauopathy link proposed in this study. As noted above, this regional finding derives from an uncorrected subfield-by-subfield comparison and should be interpreted as hypothesis-generating rather than confirmatory.

Beyond this global enrichment pattern, examination of the individual DEP dataset revealed that CSNK2A1 was significantly overexpressed in the CA3 region of patients with AD relative to controls. CSNK2A1 was prioritized earlier in this study based on network centrality and its dual, though incompletely characterized, role as a tau-associated kinase and negative regulator of autophagy. This differential protein abundance is consistent with, though not direct evidence for, an altered regulatory role for CSNK2A1 in tauopathy pathogenesis, since protein-level changes alone do not establish whether its autophagy-related function is correspondingly affected. Taken together with the exploratory nature of the regional comparison noted above, this finding should be regarded as a correlative signal that complements, rather than independently confirms, the network-based prioritization of CSNK2A1 in this study.

## Discussion

Tauopathies represent one of the greatest therapeutic challenges in neurology due to the complexity of the mechanisms regulating disease progression (Cummings et al., 2023). The accumulation of hyperphosphorylated tau (Reddy and Oliver, 2019), impaired proteostasis (Barmaki et al., 2023), mitochondrial dysfunction (Shi et al., 2025), and autophagy deterioration (Samimi et al., 2020) are part of an interconnected network, which explains the limited success of single-target strategies (Bi et al., 2025). In this context, we developed a computational method to rationally prioritize fungal metabolites that could modulate multiple regulatory nodes that link autophagy and tau pathology (Bonetto et al., 2025).

The main contribution of our study lies not only in the identification of promising metabolites, but also in the development of a hierarchical workflow that combines complementary computational methodologies to rationally reduce a vast chemical space to a limited number of candidates with high biological plausibility (Joshi et al., 2025). First, network pharmacology enabled the identification of proteins relevant to modulating autophagy in the context of tauopathies. Subsequently, molecular docking, molecular dynamics simulations, and MM/GBSA rescoring enabled the evaluation of the structural feasibility and stability of protein–ligand interactions (Alonso et al., 2006; Genheden and Ryde, 2015). Next, QSAR modeling provided a ligand-based estimate of the expected binding affinity for the candidates included within the applicability domain of each model (Cherkasov et al., 2014). Finally, the integration of human proteomic data enabled the contextualization of the predictions within a clinically relevant scenario. The consistent identification of the same molecular targets and candidate metabolites through structure-based methodologies substantially increases confidence in the results. In addition, QSAR and proteomic analyses provided additional and partially convergent evidence whenever their respective applicability domains allowed it. This convergence across independent approaches reduces the likelihood of false positives associated with any individual method (Moshawih et al., 2024). Importantly, the prioritization process did not begin with target-based analyses, but with a systematic evaluation of the physicochemical properties of the fungal metabolite library. Initial filtering based on predicted blood–brain barrier permeability and drug-likeness substantially reduced the chemical space and enriched the selection with molecules compatible with the development of drugs targeting the central nervous system.

Our computational approach identified PPARG, GSK3β, and CSNK2A1, three regulators that connect autophagy with key processes involved in the progression of tauopathies. We selected these three targets based on their established mechanistic relevance to tau pathology and autophagy regulation, rather than on network centrality alone. Notably, these are upstream regulators integrating energy metabolism, stress response, and tau phosphorylation, rather than structural components of the core autophagic machinery, which were not represented among the overlapping targets. This finding suggests that modulation of these proteins could simultaneously influence several mechanisms involved in the disease. This mechanistic diversity, detailed in the Introduction, supports the biological relevance of the targets prioritized by our computational strategy and strengthens their potential as multi-target therapeutic candidates for tauopathies.

An important aspect of the workflow was that molecular docking alone was not used as the final selection criterion. Instead, docking predictions were further challenged by long-timescale molecular dynamics simulations and MM/GBSA binding free energy calculations, allowing compounds with apparently favorable docking scores but unstable binding modes to be discarded. For example, the top-ranked docking hits Ergovaline and Lysergic Acid dissociated from the binding pocket during molecular dynamics simulations and were therefore replaced by compounds showing greater structural stability. This is consistent with the reduction in false positives achieved by integrating complementary methods, as noted above.

Emodin served as a valuable reference compound throughout the computational pipeline because its previously reported activity against PPARG, GSK3β, and CSNK2A1 enabled benchmarking of the different in silico approaches(Yamada et al., 2005; Gebhardt et al., 2010; (Yamada et al., 2005; Gebhardt et al., 2010b; Li et al., 2025b). Although several fungal metabolites outperformed emodin in MM/GBSA binding free energy calculations, emodin consistently displayed stable binding across docking, molecular dynamics simulations, and binding free energy analyses for all three targets, consistent with the convergence described above. Its QSAR-predicted affinity for GSK3β, however, falls outside the model’s applicability domain and should therefore be interpreted with lower confidence; this does not affect the independent support provided by the structural and physics-based analyses, which converge on emodin as a stable ligand for all three targets. Importantly, the superior MM/GBSA performance of compounds such as Corallocin A, Chaetopyranin, and Hericioic Acid D suggests that these metabolites may represent even more promising candidates for experimental validation than the reference compound. Furthermore, numerous experimental studies have shown that emodin exerts neuroprotective, antioxidant, and anti-inflammatory effects, modulates pathways related to AMPK and mTOR, and promotes the clearance of aggregated proteins (Wu et al., 2022b; Saha and Ahmad, 2024). Although these precedents do not directly validate the proposed interactions, they provide biological plausibility to our predictions.

Beyond emodin, metabolites belonging to structurally very different families, such as erinacerins, corallocin A, and ergocryptine, also successively passed the various selection criteria, indicating that the modulation of autophagy can be achieved through multiple independent chemical solutions. This expands the possibilities for optimization via medicinal chemistry and reduces dependence on a single lead compound.

From a translational perspective, *Hericium erinaceus* deserves special consideration. Unlike most of the fungi evaluated, it is a species widely cultivated for food and medicinal purposes, for which established production protocols via industrial-scale fermentation exist (Gonkhom et al., 2021). This characteristic could facilitate the reproducible procurement of bioactive metabolites and accelerate future preclinical studies. Conversely, other species included in this work present greater regulatory or production challenges; however, this does not diminish the interest in their metabolites as lead structures for chemical optimization programs, particularly given recent advances in metabolic engineering, precision fermentation, and heterologous reconstruction of biosynthetic pathways that offer alternatives for producing metabolites of interest regardless of the original organism (Harvey et al., 2018).

Incorporating human brain tissue proteomic data provided an independent, patient-derived layer of biological context that most network pharmacology studies lack - most conclude with target identification or structural evaluation of protein-ligand interactions, without determining whether those targets show evidence of alteration in human disease. This analysis supported the overall association between autophagy dysregulation and tauopathy identified by the computational pipeline, with enrichment of autophagy-and lysosome-related pathways detected specifically in the CA3 hippocampal subfield. As this regional comparison was exploratory and not corrected for the number of subfields tested (see Methods), this finding is best interpreted as raising the possibility of subfield-specific vulnerability rather than establishing it, and warrants replication in independent cohorts before being taken as evidence that autophagy impairment is non-uniformly distributed across the hippocampus.

In the analyzed cohort, CSNK2A1 was the only one of the three molecular targets whose protein abundance was significantly different between patients and controls. However, it is important to consider that a single differential-abundance finding from one proteomic cohort and one hippocampal subfield, indicates that CSNK2A1 levels are altered in AD tissue, but does not by itself establish that its autophagy-regulatory function is impaired. This finding is consistent with existing evidence identifying CK2 as a regulator of protein homeostasis across multiple pathological contexts, including a proposed role in tau hyperphosphorylation via the SET/PP2A axis in AD, although the overall evidence for CK2 as a therapeutic target in neurodegeneration remains preliminary (Borgo et al., 2021). The absence of significant changes for PPARG and GSK3β does not necessarily imply lower biological relevance, since both proteins are subject to post-translational regulation through phosphorylation, interaction with cofactors, ligand binding, and changes in their subcellular localization, mechanisms that can modify their activity without altering their expression levels (Suskiewicz, 2024).

The main limitation of the present work is its completely in silico nature. All tools employed constitute prioritization and hypothesis-generation methodologies, but none alone demonstrates the existence of a functional molecular interaction or the biological efficacy of the identified compounds. Consequently, the prioritized metabolites must be interpreted as candidates for experimental validation and not as potentially validated therapeutic agents. However, this limitation is inherent to the initial stages of computational drug discovery and does not represent a specific weakness of the developed pipeline. In this sense, the next steps must include the biophysical validation of protein-ligand interactions, the evaluation of the metabolites’ capacity to modulate autophagic flux in cellular models, the analysis of their effects on tau accumulation and toxicity, and subsequently, their characterization in animal models of tauopathy. Likewise, the exploration of structure-activity relationships and the chemical optimization of the identified compounds could improve their pharmacological properties and favor their preclinical development.

## Conclusion

This study presents an integrated computational pipeline that combines network pharmacology, structure-based modelling, ligand-based prediction, and human proteomic data to prioritize fungal-derived, multi-target autophagy modulators for tauopathies. PPARG, GSK3B, and CSNK2A1 emerged as biologically plausible targets, and six fungal metabolites were consistently supported by convergent structural evidence, with QSAR and proteomic analyses providing additional, partial support for a subset of these candidates. As all evidence generated here is computational or correlative, these metabolites should be regarded as candidates for experimental validation rather than confirmed therapeutic agents, and the next steps outlined above will be essential to establish their biological activity. Beyond these specific compounds, this work provides a prioritization framework that is extensible to other complex diseases.

## Author Contributions

Angel Ramon Torres Mc Cook: Methodology, Software, Investigation, Formal Analysis, Data Curation, Visualization, Writing – original draft. Camila Mimura: Writing & editing. Lautaro Damian Alvarez: Methodology, Software, Investigation, Data Curation, Visualization, Supervision, Writing. Ana Clara Liberman: Conceptualization, Supervision, Writing. All authors contributed to the article and approved the submitted version.

---

**Supplementary Figure 1. Chemical structures of the 27 compounds derived from Ergot Fungi included in the compound library used in this study.**

**Supplementary Figure 2. Chemical structures of the 75 compounds derived from Lion’s Mane included in the compound library used in this study.**

**Supplementary Figure 3. Chemical structures of the 38 compounds derived from Magic Mushrooms included in the compound library used in this study.**

