## Supplementary Figures for "Network Pharmacology-Guided Discovery of Fungal Autophagy Modulators for Tauopathies: Structural and Proteomic Evidence"

### Supplementary Figure 1

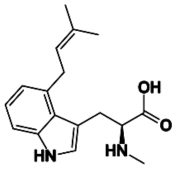
4-(3-Methylbut-2-enyl)-L-abrine

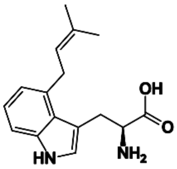
4-(3-Methylbut-2-enyl)-L-tryptophan

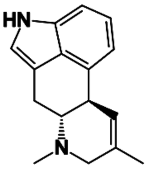
Agroclavine

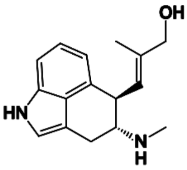
Chanoclavine I

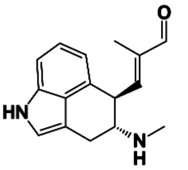
Chanoclavine I Aldehyde

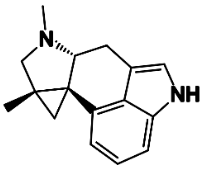
Cycloclavine

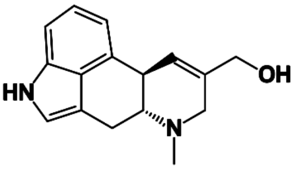
Elymoclavine

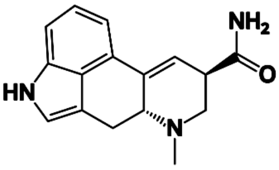
Ergine

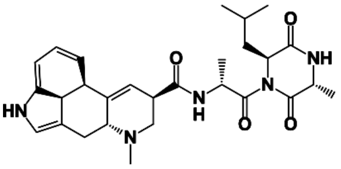
Ergobalansam

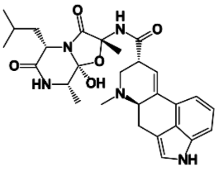
Ergobalansine

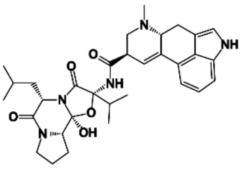
Ergocryptine

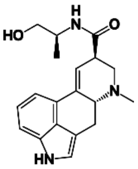
Ergonovine

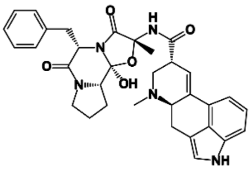
Ergotamine

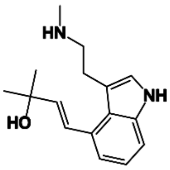
Ergotryptamine

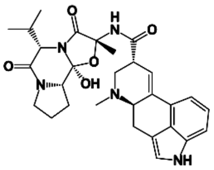
Ergovaline

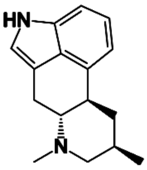
Festuclavine

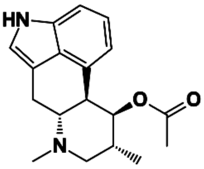
Fumigaclavine A

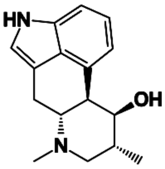
Fumigaclavine B

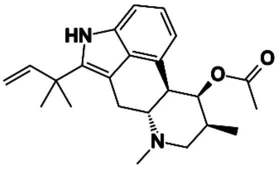
Fumigaclavine C

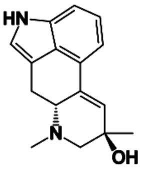
Isosetoclavine

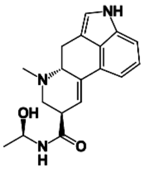
LAH

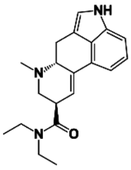
LSD

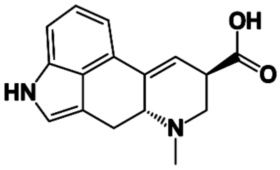
Lysergic Acid

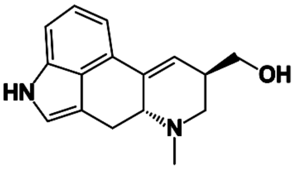
Lysergol

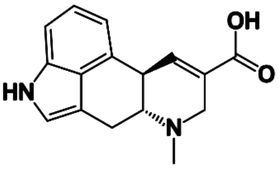
Paspalic Acid

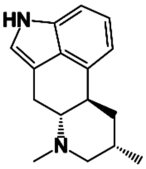
Pyroclavine

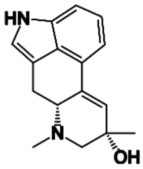
Setoclavine

### Supplementary Figure 2

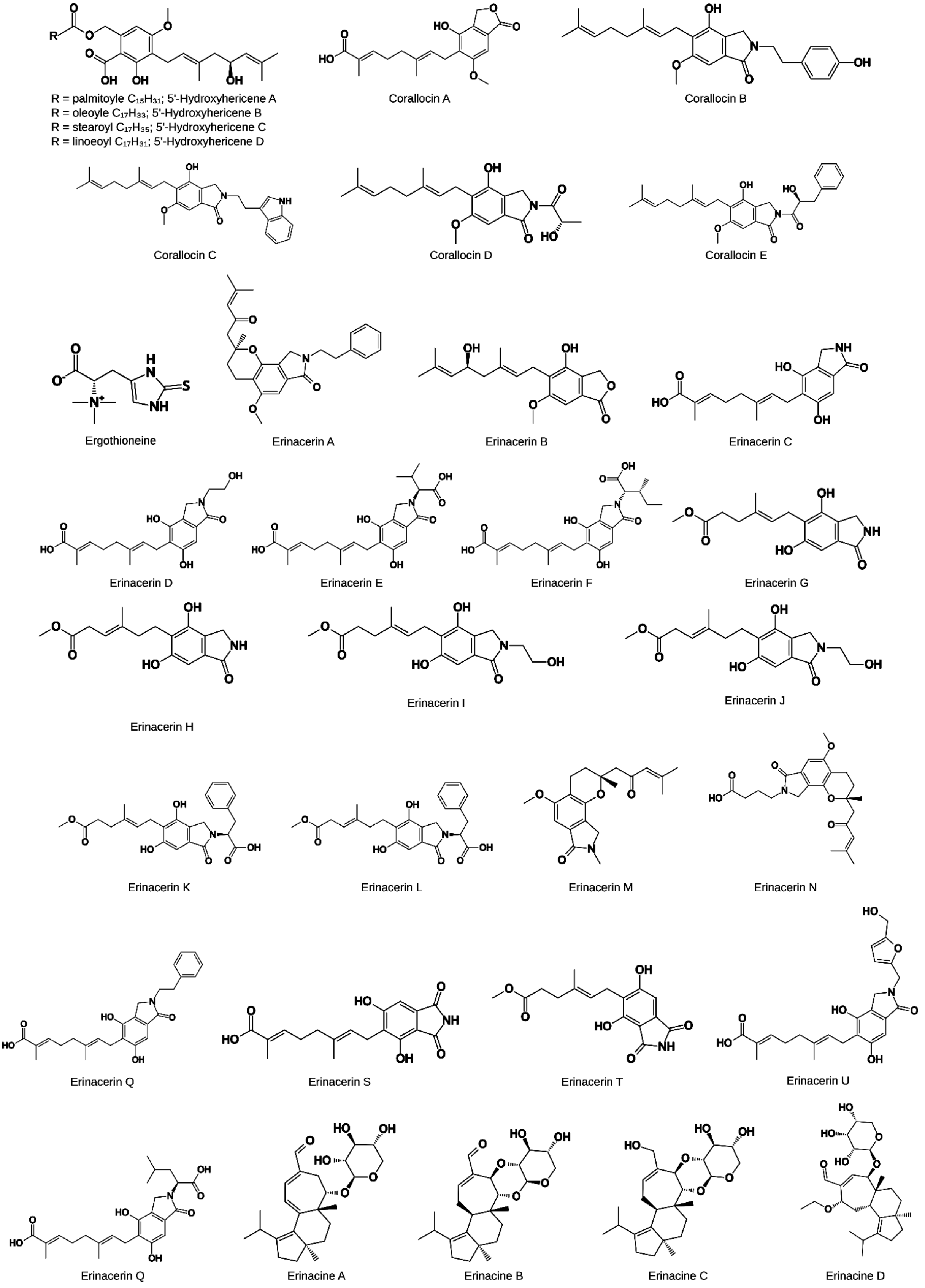

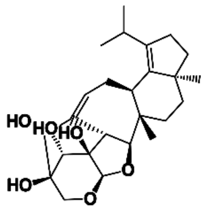
Erinacine E

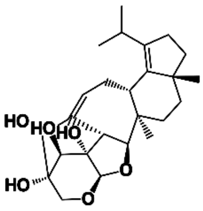
Erinacine F

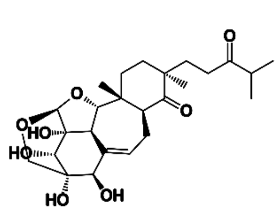
Erinacine G

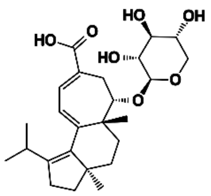
Erinacine H

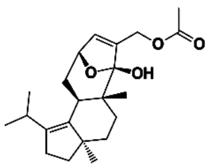
Erinacine I

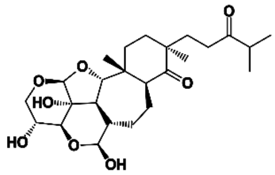
Erinacine J

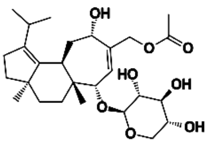
Erinacine K

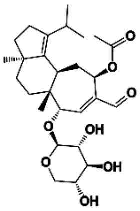
Erinacine P

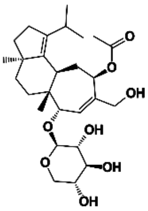
Erinacine Q

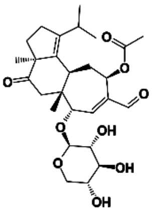
Erinacine R

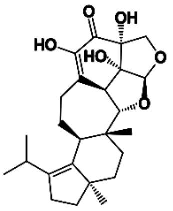
Erinacine S

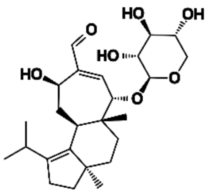
Erinacine T

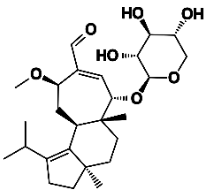
Erinacine U

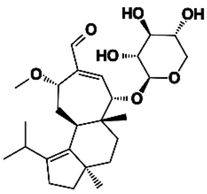
Erinacine V

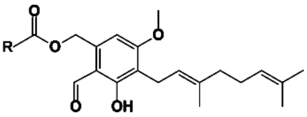
R = palmitoyle $C_{15}H_{31}$; Hericene A
R = oleoyle $C_{17}H_{33}$; Hericene B
R = stearoyl $C_{17}H_{35}$; Hericene C
R = linoeoyl $C_{17}H_{31}$; Hericene D

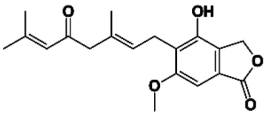
Hericenone A

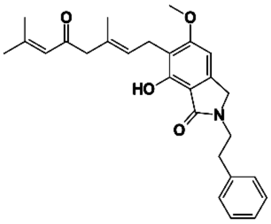
Hericenone B

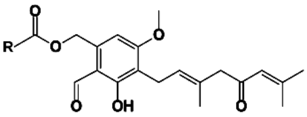
R = palmitoyle $C_{15}H_{31}$; Hericenone C
R = stearoyl $C_{17}H_{35}$; Hericenone D
R = linoeoyl $C_{17}H_{31}$; Hericenone E

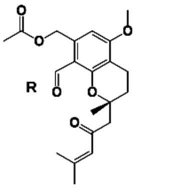
R = palmitoyle $C_{15}H_{31}$; Hericenone F
R = stearoyl $C_{17}H_{35}$; Hericenone G
R = linoeoyl $C_{17}H_{31}$; Hericenone H

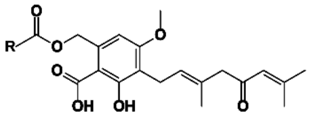
R = palmitoyle $C_{15}H_{31}$; Hericenone L

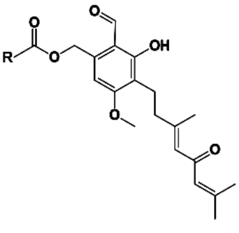
R = palmitoyle $C_{15}H_{31}$; Hericenone P
R = stearoyl $C_{17}H_{35}$; Hericenone Q
R = linoeoyl $C_{17}H_{31}$; Hericenone O

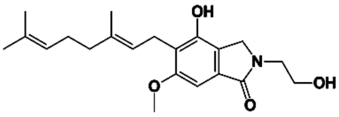
Hericerin A

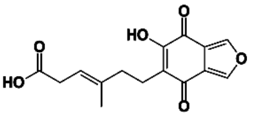
Hericiofuranoic Acid

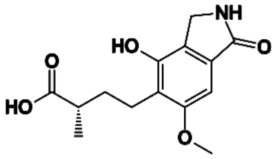
Hericioic Acid A

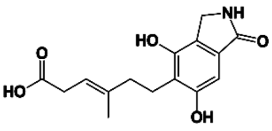
Hericioic Acid B

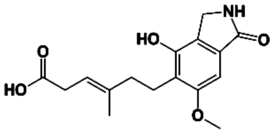
Hericioic Acid C

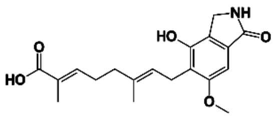
Hericioic Acid D

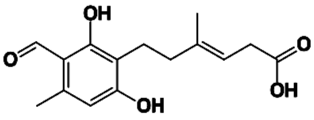
Hericioic Acid E

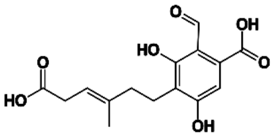
Hericioic Acid F

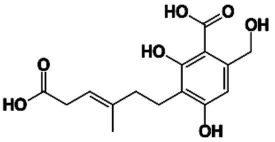
Hericioic Acid G

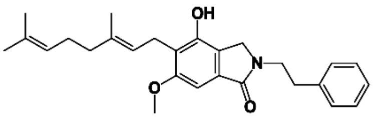
Isohericerin

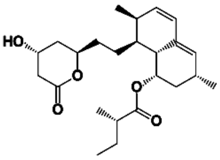
Lovastatin

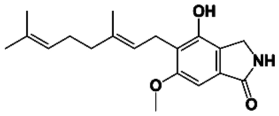
NDPIH

### Supplementary Figure 3

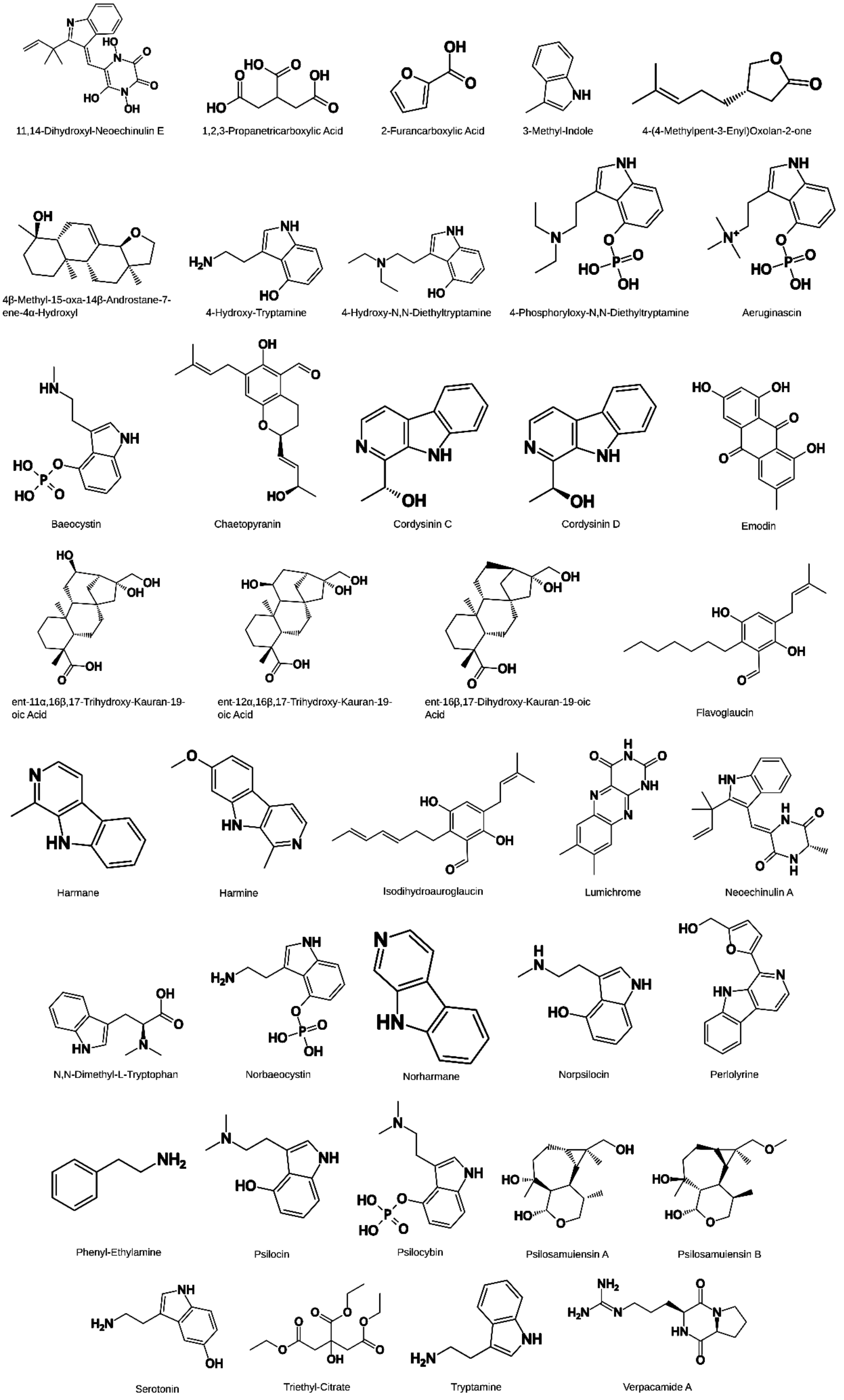
